# Colloidal aggregators in biochemical SARS-CoV-2 repurposing screens

**DOI:** 10.1101/2021.08.31.458413

**Authors:** Henry R. O’Donnell, Tia A. Tummino, Conner Bardine, Charles S. Craik, Brian K. Shoichet

**Affiliations:** Department of Pharmaceutical Chemistry, University of California San Francisco (UCSF), San Francisco, CA, USA; Graduate Program in Pharmaceutical Sciences and Pharmacogenomics, UCSF, San Francisco, CA, USA; QBI COVID-19 Research Group (QCRG), San Francisco, CA, USA; Graduate Program in Chemistry & Chemical Biology, UCSF, San Francisco, CA, USA

## Abstract

To fight the SARS-CoV-2 pandemic, much effort has been directed toward drug repurposing, testing investigational and approved drugs against several viral or human proteins *in vitro*. Here we investigate the impact of colloidal aggregation, a common artifact in early drug discovery, in these repurposing screens. We selected 56 drugs reported to be active in biochemical assays and tested them for aggregation by both dynamic light scattering and by enzyme counter screening with and without detergent; seventeen of these drugs formed colloids at concentrations similar to their literature reported IC_50_s. To investigate the occurrence of colloidal aggregators more generally in repurposing libraries, we further selected 15 drugs that had physical properties resembling known aggregators from a common repurposing library, and found that 6 of these aggregated at micromolar concentrations. An attraction of repurposing is that drugs active on one target are considered de-risked on another. This study suggests not only that many of the drugs repurposed for SARS-CoV-2 in biochemical assays are artifacts, but that, more generally, when screened at relevant concentrations, drugs can act artifactually via colloidal aggregation. Understanding the role of aggregation, and detecting its effects rapidly, will allow the community to focus on those drugs and leads that genuinely have potential for treating COVID-19.

Table of Contents Graphic

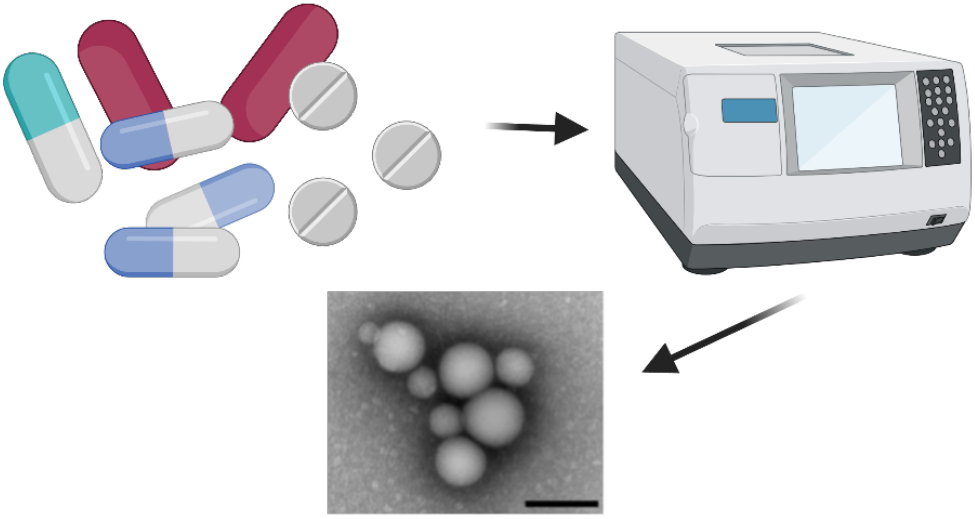

## Introduction

Drug repurposing is an attractive idea in the face of a global pandemic, when rapid antiviral drug development is crucial. While the historical pragmatism of this approach has drawn scrutiny,^1, 2^ drug repurposing has the potential to dramatically cut both the time and cost needed to develop a new therapeutic.^3^ Repurposing campaigns typically screen curated libraries of thousands of approved drugs and investigational new drugs (INDs), and several assays have been developed to test these libraries for activity against SARS-CoV-2 ^4–6^. Most high throughput, biochemical screens were developed to detect activity against two proteins that are used in viral infection and maturation: the human ACE-2 (Angiotensin converting enzyme 2) and 3CL-Pro,^7^ the major polypeptide processing protease of SARS-2-CoV-2.

When testing molecules for biochemical activity at micromolar concentrations, it is important to control for artifacts^8–12^ including colloidal aggregation, which is perhaps the single most common artifact in early drug discovery.^13, 14^ Drugs, though in many ways de-risked, are not immune to aggregation and artifactual behavior when screened at relevant concentrations ^15, 16^ (though they are not expected to aggregate at on-target relevant concentrations). Knowing this, we wondered if colloidal aggregation was causing false positives in some COVID-19 drug repurposing studies, especially since several known aggregators, such as manidipine and methylene blue, were reported as apparently potent hits for Covid-19 targets.^17, 18^

Aggregation is a is common source of false positives in early drug discovery,^19^ arising from spontaneous formation of colloidal particles when organic, drug-like molecules are introduced into aqueous media.^15, 16, 19, 20^ The resulting liquid particles are densely packed spheres^21^ that promiscuously inhibit proteins by sequestering them on the colloid surface,^22^ where they suffer partial unfolding.^23^ The resulting inhibition is reversable by disruption of the colloid, and is characterized by an incubation effect on the order of several minutes due to enzyme crowding on the surface of the particle.^24^ Colloids often can be disrupted by the addition of small amounts, often sub-critical micelle concentrations, of non-ionic detergent such as Triton-X 100.^25^ Accordingly, addition of detergent is a common perturbation to rapidly detect aggregates in counter screens against model enzymes such as AmpC ß-lactamase or malate dehydrogenase (MDH). Aggregation can be physically detected by biophysical techniques such as nuclear magnetic resonance (NMR)^26^ and by dynamic light scattering (DLS), as the colloids typically form particles in the 50 to 500 nm radius size range, which is well-suited to measurement by this latter technique.

Here we investigate the role of colloidal aggregation as a source of false positives in drug repurposing studies for SARS-CoV-2 targets. We focused on *in vitro*, ACE2 and 3CL-Pro screens since these are relevant for aggregation. We searched the literature and compiled the top hits from 12 studies^18, 27–37^ and these were then visually inspected and ordered. How the results of this study may impact the design of future repurposing screens both for SARS-2 and for other indicators, will be considered.

## Results

### Colloidal aggregators are common hits in drug repurposing screens for SARS-CoV-2

We tested 56 drugs for colloidal aggregation that had been reported to be active in biochemical repurposing screens against SARS-CoV-2^18, 27–30, 32, 38^ (**SI Table 1**). Five criteria were used to investigate whether reported hits formed colloidal aggregates: **a.** particle formation indicated by scattering intensity, **b.** clear autocorrelation curves, **c.** an MDH IC_50_ in the micromolar – high nanomolar range, **d.** restoration of MDH activity with the addition of detergent, and less stringently **e.** high Hill slopes in the inhibition concentration response curves (**Fig 1**).

**Fig 1).**
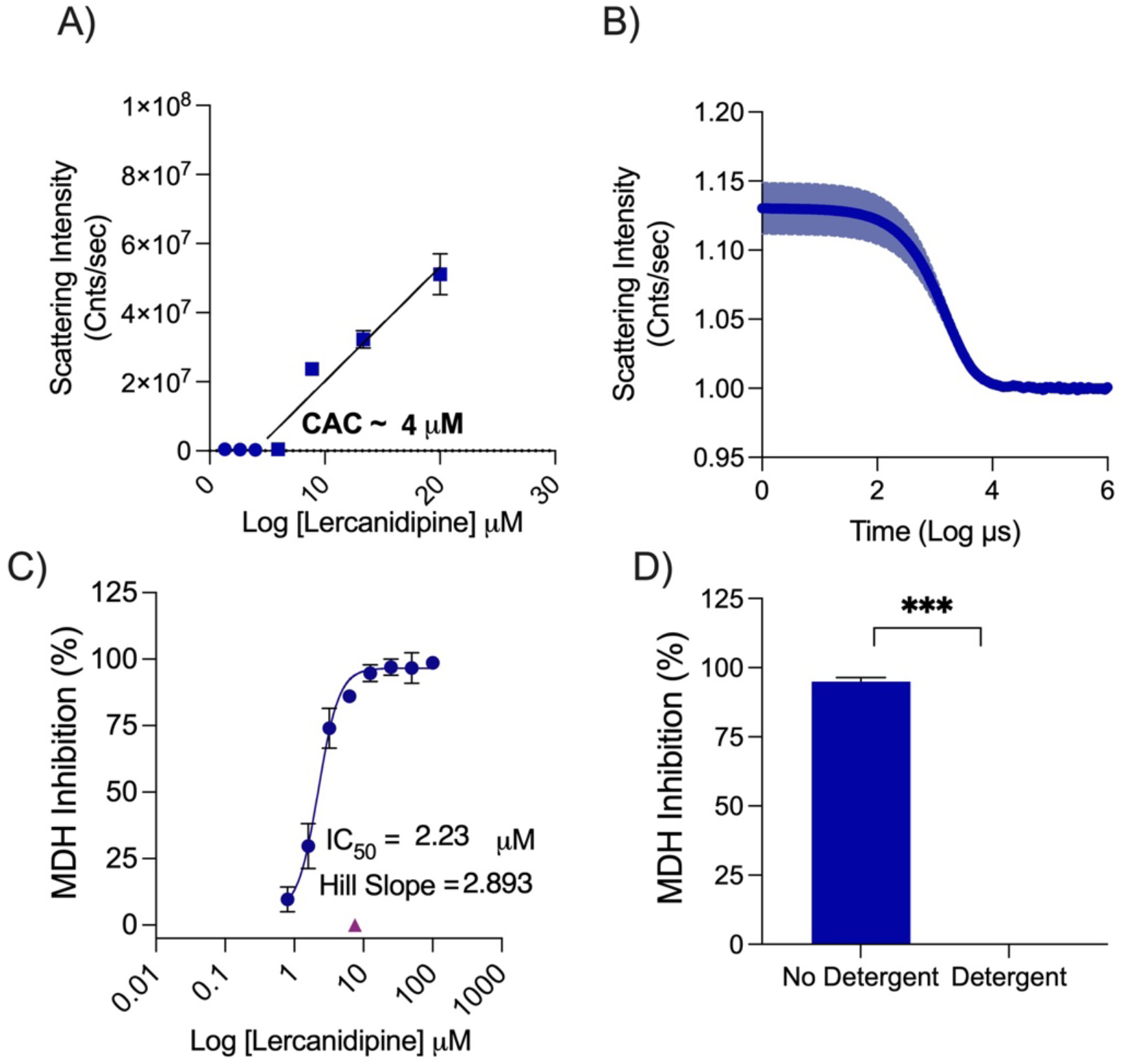
Lercanidipine’s behavior as an aggregator. (A) Critical aggregation concentration determined using scattering intensity measured on DLS. (B) Autocorrelation curve from DLS at 100 μM. (C) Dose response measured against MDH and showing the Hill slope. (D) MDH inhibition measured with or without 0.01% TritonX-100 at 7.5 μM.

Using the literature reported IC_50_ for the repurposed drugs as a starting point, we tested each drug for MDH inhibition and calculated the IC_50_ and Hill slope. We used IC_50_ values from the MDH concentration response curves and tested for detergent sensitivity at 3-fold the MDH IC_50_ (**Fig 2**). Next, we calculated the critical aggregation concentration (CAC) by measuring normalized scattering intensity on the DLS, any point above 1×10^6^ was considered from the aggregated form. By plotting a best fit line for aggregating concentrations and non-aggregating concentrations, the CAC was given by the point of intersection (**Fig 3**). We also measured the DLS auto correlation curve as a criterion: if this was well-formed it gave further confidence (**SI Fig 1**).

**Fig 2).**
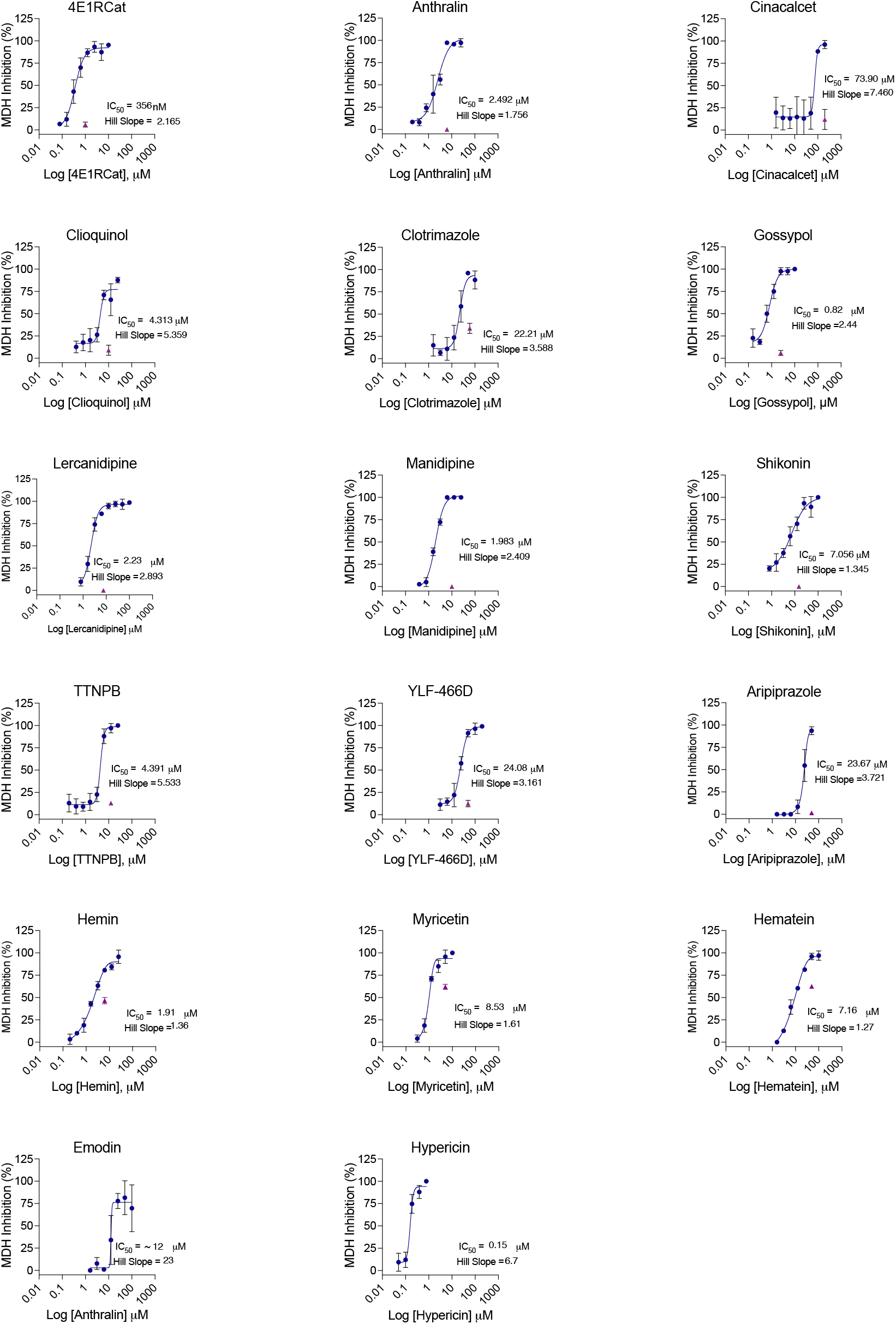
MDH inhibition concentration-response curves for literature actives. IC_50_ and Hill slopes are shown. Purple triangles indicate single point MDH inhibition with the addition of 0.01% TritonX-100, tested at 3x IC_50_. All measurements in triplicate.

**Fig 3).**
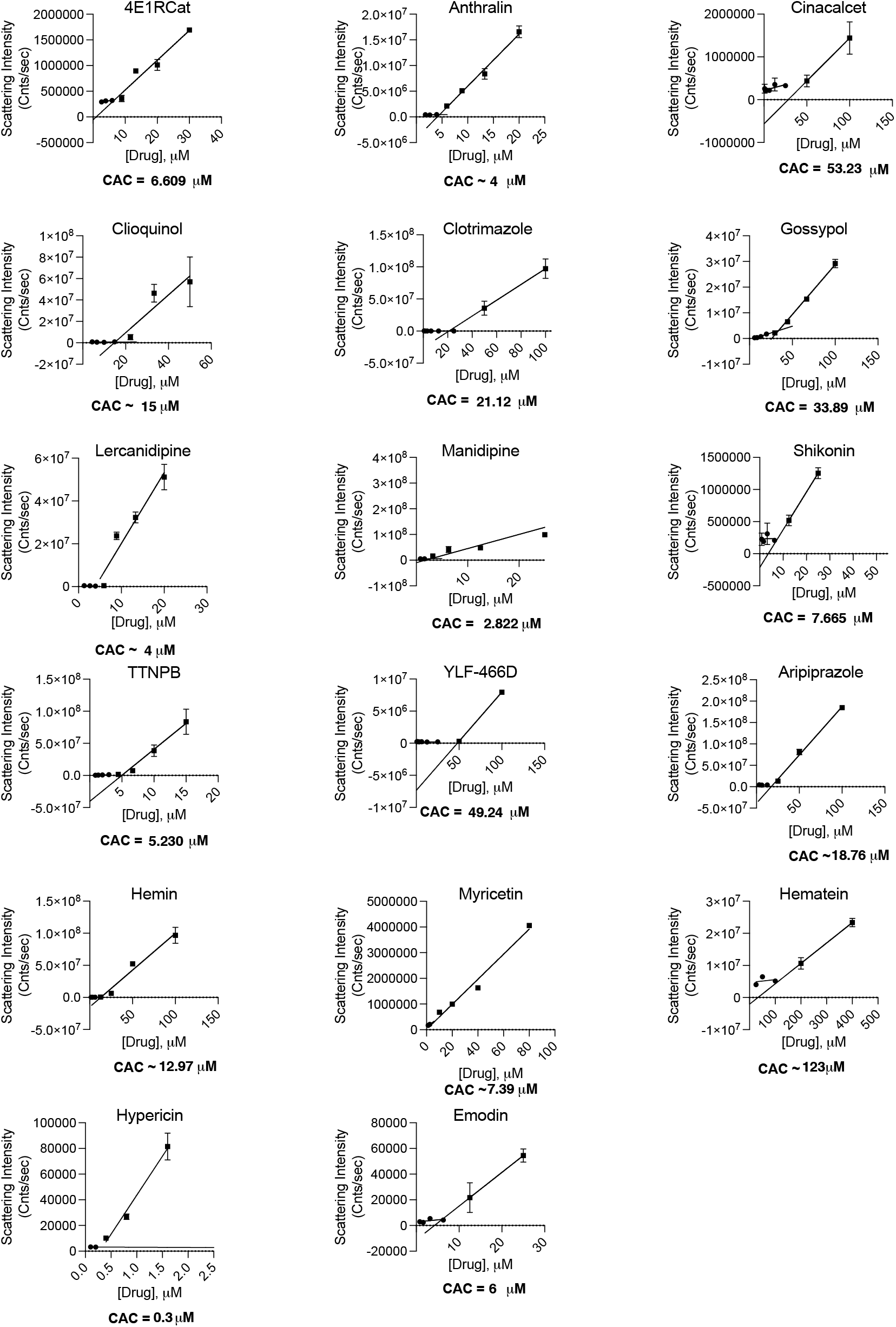
Critical aggregation concentrations for literature actives. CAC is determined by finding the intersection of two best fit lines for points with scattering intensity above or below 1×10^6^. All measurements in triplicate.

Seventeen molecules formed well-behaved particles detectable by DLS with clean autocorrelation curves, and inhibited MDH in the absence of, but not the presence of 0.01% Triton X-100; these seem to be clear colloidal aggregators (**Table 1, Fig 2, Fig 3**). Both DLS-based critical aggregation concentrations and MDH IC_50_ values were in the range of the primary IC_50_s reported in the literature against the two SARS-CoV-2 enzymes; indeed, molecules like Gossypol, manidipine, and TTNPB inhibited the counterscreening enzyme MDH even more potently than they did either ACE2 or 3CL-Pro. For most of the 17 drugs, the Hill slopes were high, though for several clear aggregators, such as Hemin and Shikonin, they were only in the 1.3 – 1.4 range. Hill slope depends on the enzyme concentration to true KD ratio and so will vary from assay to assay ^39^. While many consider it a harbinger of aggregation, we take it as only a soft criterion.

**Table 1.**
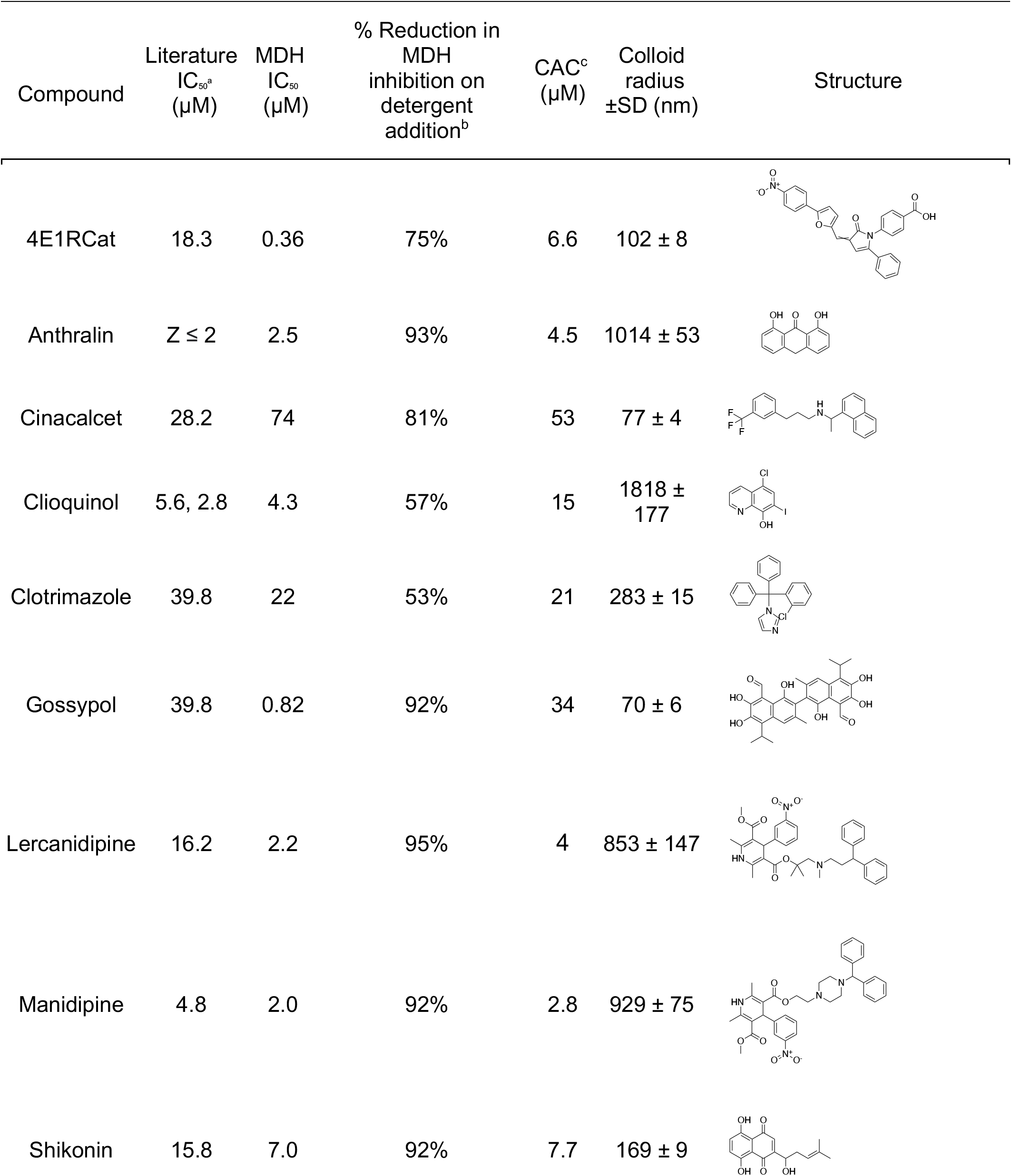
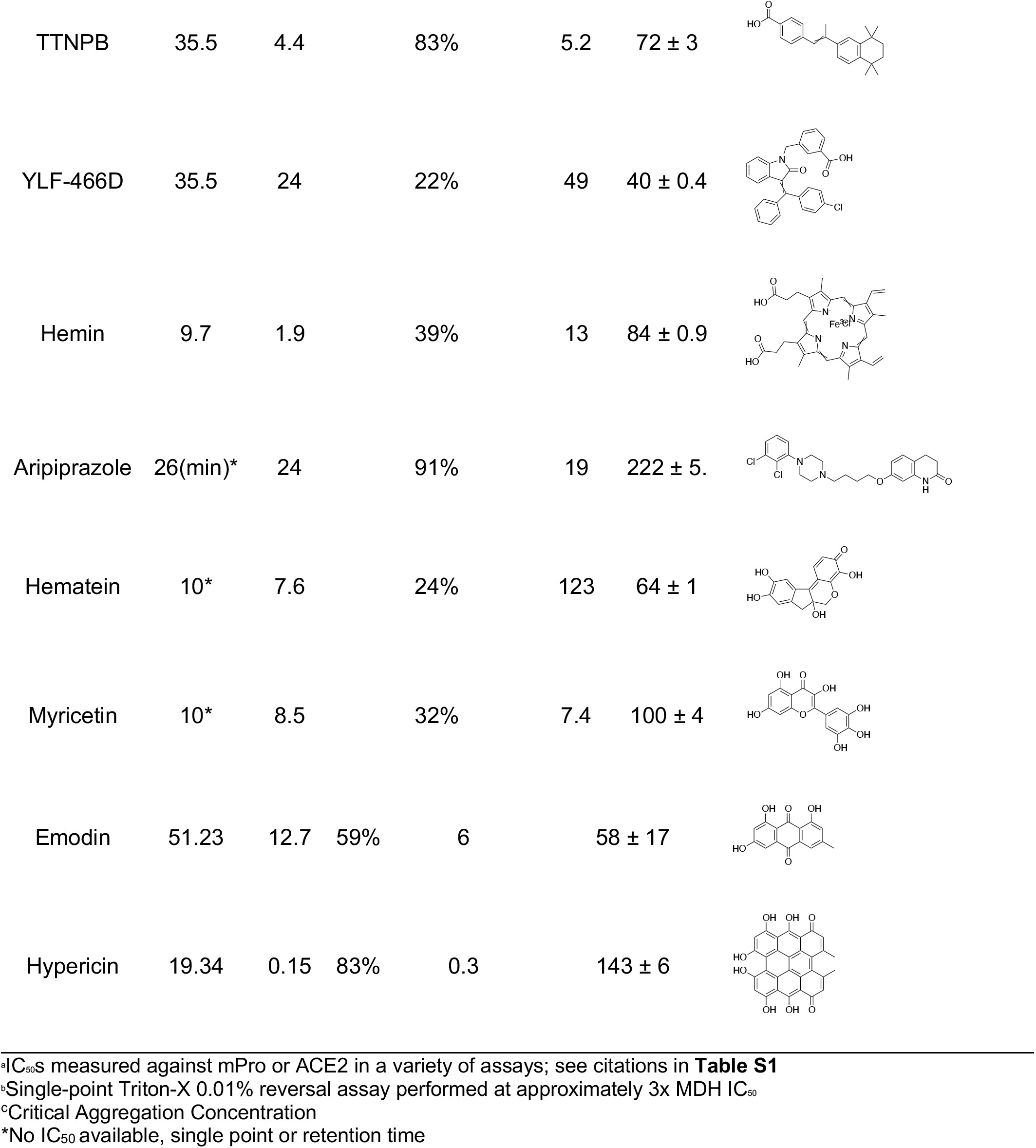
Literature SARS-CoV-2 repurposing hits shown to cause colloidal aggregation.

| Compound | Literature<br>IC<sub>50</sub><sup>a</sup><br>(μM) | MDH<br>IC<sub>50</sub><br>(μM) | % Reduction in<br>MDH<br>inhibition on<br>detergent<br>addition<sup>b</sup> | CAC<sup>c</sup><br>(μM) | Colloid<br>radius<br>±SD (nm) | Structure |
| --- | --- | --- | --- | --- | --- | --- |
| 4E1RCat | 18.3 | 0.36 | 75% | 6.6 | 102 ± 8 |  |
| Anthralin | Z ≤ 2 | 2.5 | 93% | 4.5 | 1014 ± 53 |  |
| Cinacalcet | 28.2 | 74 | 81% | 53 | 77 ± 4 |  |
| Clioquinol | 5.6, 2.8 | 4.3 | 57% | 15 | 1818 ± 177 |  |
| Clotrimazole | 39.8 | 22 | 53% | 21 | 283 ± 15 |  |
| Gossypol | 39.8 | 0.82 | 92% | 34 | 70 ± 6 |  |
| Lercanidipine | 16.2 | 2.2 | 95% | 4 | 853 ± 147 |  |
| Manidipine | 4.8 | 2.0 | 92% | 2.8 | 929 ± 75 |  |
| Shikonin | 15.8 | 7.0 | 92% | 7.7 | 169 ± 9 |  |
| TTNPB | 35.5 | 4.4 | 83% | 5.2 | 72 ± 3 |  |
| YLF-466D | 35.5 | 24 | 22% | 49 | 40 ± 0.4 |  |
| Hemin | 9.7 | 1.9 | 39% | 13 | 84 ± 0.9 |  |
| Aripiprazole | 26(min)* | 24 | 91% | 19 | 222 ± 5. |  |
| Hematein | 10* | 7.6 | 24% | 123 | 64 ± 1 |  |
| Myricetin | 10* | 8.5 | 32% | 7.4 | 100 ± 4 |  |
| Emodin | 51.23 | 12.7 | 59% | 6 | 58 ± 17 |  |
| Hypericin | 19.34 | 0.15 | 83% | 0.3 | 143 ± 6 |  |

<sup>a</sup>IC<sub>50</sub>s measured against mPro or ACE2 in a variety of assays; see citations in **Table S1**
<sup>b</sup>Single-point Triton-X 0.01% reversal assay performed at approximately 3x MDH IC<sub>50</sub>
<sup>c</sup>Critical Aggregation Concentration
\*No IC<sub>50</sub> available, single point or retention time

A characteristic example of a reported hit that is likely acting artifactually through colloidal aggregation is the calcium channel blocker lercanidipine, which has been reported to inhibit 3CL-Pro with an IC_50_ of 16.2 μM ^18^. Lercanidipine satisfies our five criteria for aggregation: in aqueous buffer if forms particles that can be detected by a 10-fold increase in DLS scattering intensity (Cnts/sec); by a clearly defined autocorrelation curve in the DLS; it inhibits the counter-screening enzyme MDH with an IC_50_ of 2.2 μM, while MDH activity is restored on addition of 0.01% Triton-X 100 detergent (**Fig 1**). In the absence of detergent, lercanidipine inhibits MDH with a Hill slope of 2.9.

In addition to the 17 molecules that passed all five criteria for aggregation, another 19 molecules were more ambiguous, either forming particles by DLS but not inhibiting MDH, or inhibiting MDH in a detergent-dependent manner but not forming particles detectable by DLS (**SI Table 1)**. These 19 drugs may also be acting artifactually, however further investigation is needed to determine their exact mechanisms. For this study, we focused only on clear colloidal aggregators.

### Molecules repurposed for 3CL-Pro show little activity against that enzyme in the presence of detergent

In addition to testing the repurposed molecules against a counter-screening enzyme like MDH, we also tested the ten that had been repurposed against 3CL-Pro against that enzyme itself. Because 3CL-Pro is unstable in buffer without either the presence of detergent or substantial amounts of serum albumin—both of which disrupt colloids ^40–42^—we could not investigate the impact of detergent with 3CL-Pro as we could do with MDH. Still, we could ask whether the drugs repurposed for 3CL-Pro inhibited the enzyme in the presence of the 0.05% Tween-20 used to keep the enzyme stable. Of the 12 drugs tested, only two had detectable potency below 200 μM in the presence of detergent, and for one of these two, 4E1RCat, their inhibition was reduced 5-fold over their literature values (18.28 to 100μM) (**Table 3, SI Figure 3**). Only hemin continued to inhibit 3CL-Pro substantially, with an IC_50_ of 25 μM (but even this was 2.6-fold less potent than its literature value). As hemin’s inhibition of MDH was disrupted by detergent (**Table 2**) and it formed clear particles by DLS (**Figure 3**, **SI Figure 3**), we further tested it against the model counter-screening enzyme AmpC β- lactamase. Hemin inhibited AmpC with an IC_50_ of 23 μM; at 25 μM hemin, addition of 0.01% (v/v) Triton X-100 fully restored enzyme activity—inhibition was abolished. Taken together, these observations further support the aggregation-based activity of these 12 repurposed drugs.

**Table 2.**
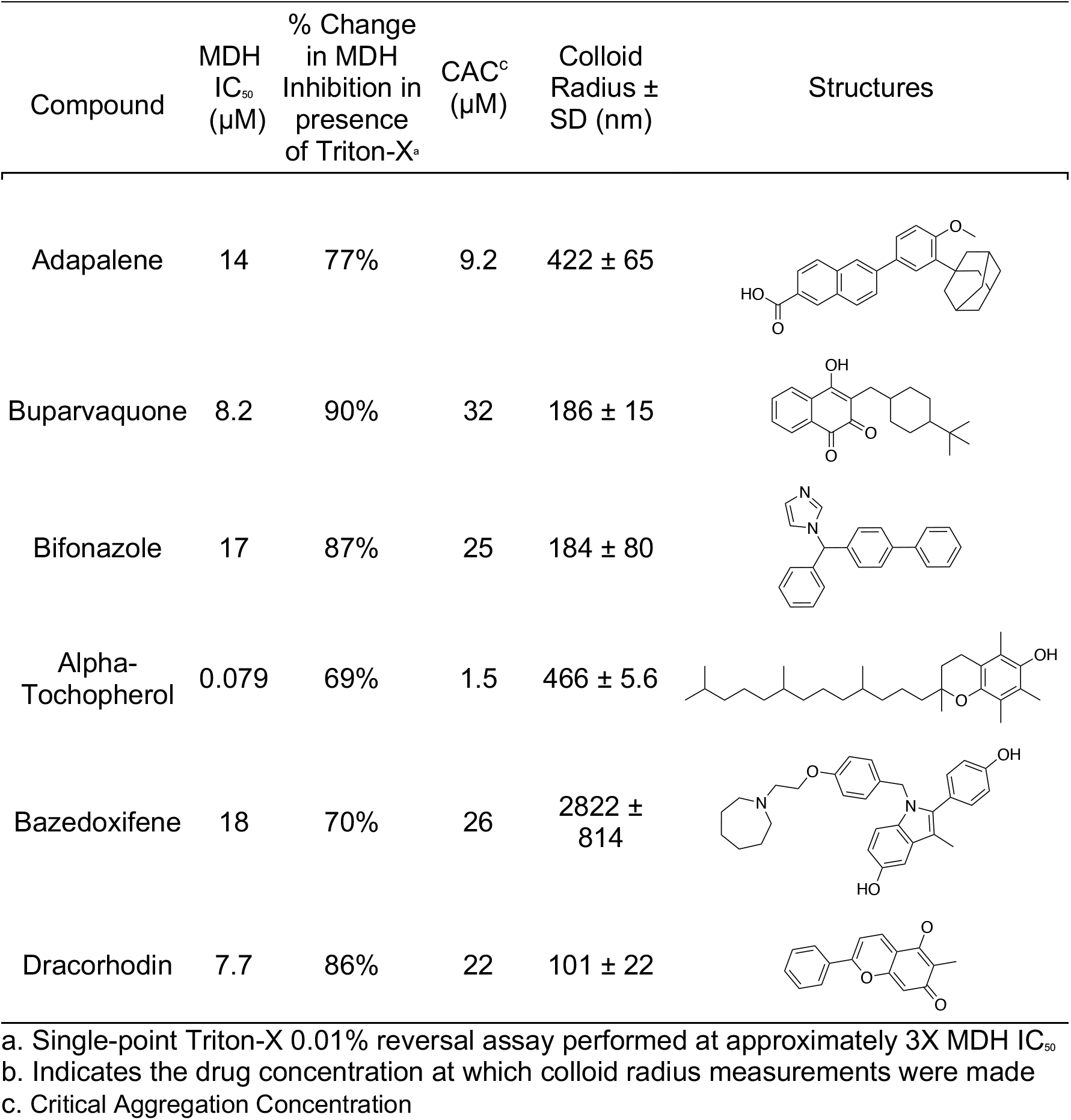
Six drugs from a repurposing library aggregate at screening-relevant concentrations.

| Compound | MDH IC<sub>50</sub> (μM) | % Change in MDH Inhibition in presence of Triton-X<sup>a</sup> | CAC<sup>c</sup> (μM) | Colloid Radius ± SD (nm) | Structures |
| --- | --- | --- | --- | --- | --- |
| Adapalene | 14 | 77% | 9.2 | 422 ± 65 |  |
| Buparvaquone | 8.2 | 90% | 32 | 186 ± 15 |  |
| Bifonazole | 17 | 87% | 25 | 184 ± 80 |  |
| Alpha-Tochopherol | 0.079 | 69% | 1.5 | 466 ± 5.6 |  |
| Bazedoxifene | 18 | 70% | 26 | 2822 ± 814 |  |
| Dracorhodin | 7.7 | 86% | 22 | 101 ± 22 |  |

a. Single-point Triton-X 0.01% reversal assay performed at approximately 3X MDH IC<sub>50</sub>
b. Indicates the drug concentration at which colloid radius measurements were made
c. Critical Aggregation Concentration

**Table 3.** Literature repurposing hits do not potently inhibit 3CL-Pro in the presence of detergent.

| Compound | Literature IC<sub>50</sub>(μM) | 3CL-Pro IC<sub>50</sub> with 0.05% Tween-20 IC<sub>50</sub>(μM) |
| --- | --- | --- |
| 4E1RCat | 18.3 | ~100 |
| Anthralin | $Z \leq 2$ | >200 |
| Clotrimazole | 39.8 | >200 |
| Gossypol | 39.8 | >200 |
| Lercanidipine | 16.2 | >200 |
| Manidipine | 4.8 | >100* |
| Shikonin | 15.8 | ~200 |
| TTNPB | 35.5 | >200 |
| YLF-466D | 35.5 | >200 |
| Hemin | 9.7 | 25 |
| Hematein | 10* | >200 |
| Emodin | 51.2 | >200 |

\*100 μM was the highest concentration used for Manidipine, instead of 200 μM

### Colloidal aggregators in repurposing libraries

Target-based drug repurposing screens are common not only for SARS-CoV-2, but for many other viruses and indeed other indications. We thought it interesting to explore, if only preliminarily, the occurrence of colloidal aggregators in drug repurposing libraries. We prioritized drugs in the widely used SelleckChem FDA-approved library as potential aggregators, using a simple chemoinformatics approach ^43^. Of the 2,336 unique drugs in the library, 73 are already known aggregators, and another 356 (16%) closely resemble known aggregators. We selected 15 of the latter for aggregation: six of these drugs satisfied our five criteria for aggregation, they inhibited MDH in the absence of, but not the presence of 0.01% Triton X-100 (**Fig 4**) and formed well-behaved particles detectable by DLS (**Fig 5**) with clean autocorrelation curves (**SI Fig 2**), often with steep Hill slopes. In aggregate, these data suggest that these six drugs are prone to colloidal aggregation at screening-relevant concentrations (**Table 2**).

**Fig 4).**
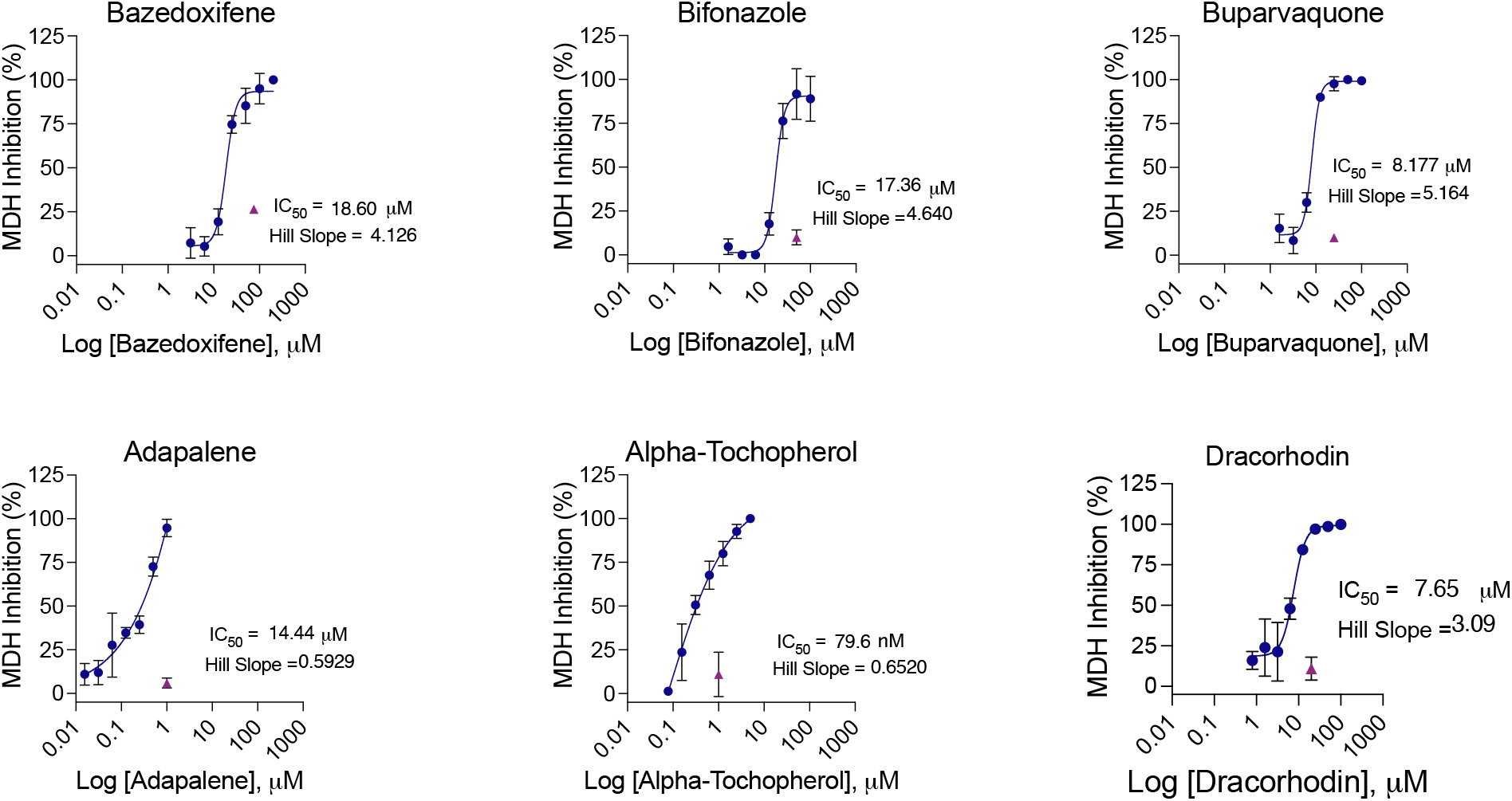
MDH inhibition dose response curves for drugs drawn from a repurposing library. All measurements were in triplicate.

**Fig 5).**
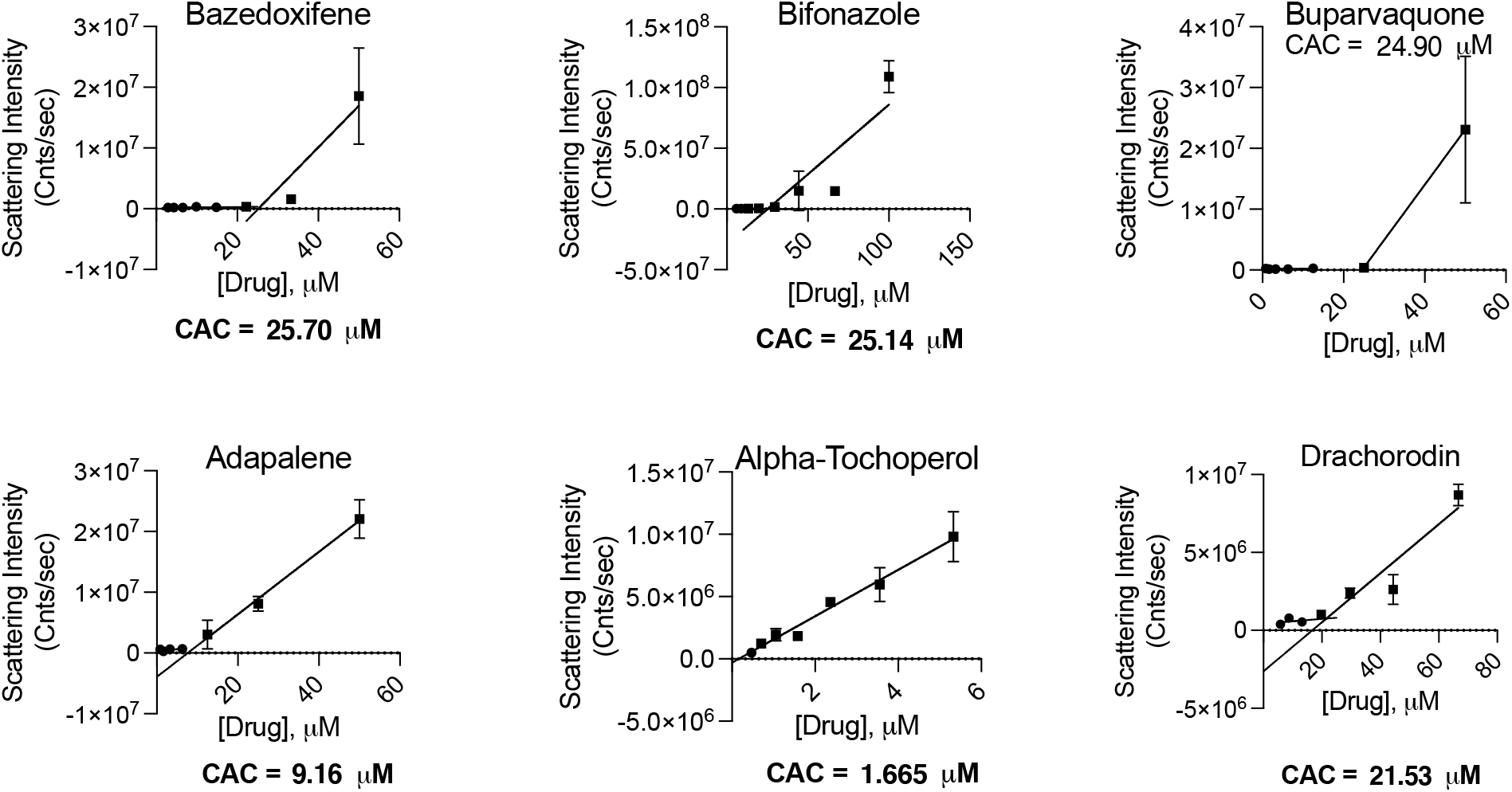
Critical aggregation concentrations for drugs drawn from a repurposing library. The CAC is determined by the intersection of two best fit lines, for points with scattering intensity above or below 1×10^6^. All measurements in triplicate.

## Discussion and Conclusions

Two broad observations from this study merit emphasis. **First**, many drugs repurposed for Covid-19 aggregate and inhibit counter-screening enzymes promiscuously at concentrations relevant to their reported IC_50_s against the Covid-19 targets (ACE2 and 3CL-Pro). Of the 56 drugs tested, 17 fulfilled all five of our criteria for acting via colloidal aggregation: **i.** they formed particles that scattered strongly by DLS with **ii.** well-behaved autocorrelation curves, **iii.** they inhibited the counter-screening enzyme malate dehydrogenase—unrelated to either ACE2 or 3CL-Pro—at relevant concentrations in the absence but **iv.** not the presence of detergent and **v.** they typically inhibited with steep Hill slopes. Each of these criteria individually is a harbinger of colloidal aggregation; combined they strongly support its occurrence. Another 19 of the 56 drugs fulfilled only some of these criteria—for instance, forming particles at relevant concentrations but not inhibiting MDH in a detergent-dependent manner. Some of these 19 may also be aggregators, while others, like those that inhibit MDH but cannot be reversed by detergent, like tannic acid, may be acting via other PAINS-like artifacts. A **second** observation from this study is that these artifacts are not so much a feature of SARS-CoV-2 repurposing, but rather reflect the behavior of drugs at screening relevant concentrations. Thus, six of fifteen drugs investigated from a general purposing library were also aggregators at micromolar concentrations. An attraction of drug repurposing is that the molecules are thought to be de-risked from the pathologies of early discovery. But at micromolar concentrations, drugs, which are often larger and more hydrophobic than the lead-like molecules found in most high-throughput screening and virtual libraries, are if anything *more* likely to aggregate, something earlier studies also support ^15, 16^.

Certain caveats should be mentioned. We do not pretend to have undertaken a comprehensive study of the increasingly large literature around drug repurposing for Covid-19. The molecules tested here represent only a subset of those investigated, drawn from an analysis of some of the literature then available. Also, we have not demonstrated that aggregation is actually occurring in the ACE-2 assay itself, though the lack of inhibition of 3CL-Pro in the presence of detergent fortifies our conclusions for the 12 molecules that inhibited this enzyme. Finally, it is important to note that just because some repurposed drugs aggregate at micromolar concentrations, the repurposing enterprise is not sunk. There are, after all, examples of drugs successfully repurposed, even for Covid-19, and some have even begun from screening hits (though typically they are subsequently modified chemically ^44^).

These caveats should not obscure the main observations from this study. Many drugs repurposed for Covid-19 in biochemical assays are aggregators—still others may be inhibiting through other artifactual mechanisms—and their promise as leads for treating the disease merits reconsideration. More broadly, drugs in repurposing libraries, though de-risked for whole body toxicity, pharmacokinetic exposure and metabolism, are not de-risked for artifactual activity at screening relevant concentrations. More encouragingly, what this study illuminates is a series of rapid assays that can rapidly distinguish drugs acting artifactually via colloidal aggregation from those drugs with true promise for treating SARS-CoV-2, and from pandemics yet to be faced.

## Experimental Section

### Literature search and chemoinformatic selection of potential aggregators

We used two approaches to identify drugs with the potential to form colloidal aggregates from repurposing screens: 1) literature searches of published SARS-CoV-2 biochemical drug screening papers including a chemoinformatic analysis of the NCATS COVID-19 OpenData Portal^37^ 3CL-Pro and ACE2 biochemical drug screens, and 2) chemoinformatic predictions of potential aggregators found in the SelleckChem FDA-approved drug library using the Aggregation Advisor tool ^43^. Literature-based keyword searches were performed using variations of the keywords *SARS-CoV-2* AND *drug repurposing* OR *drug screen*. Active molecules in papers that performed non cell-based *in vitro* drug-repurposing screens were visually inspected and prioritized for testing if they were highly hydrophobic or aromatic. Next, data from the NCATS COVID-19 OpenData Portal ^37^ drug-repurposing screens for modulators of 3CL-Pro and ACE2 activity were retrieved (accessed on 2020-09-28). In total, 12,262 and 3,405 annotations were found for compounds screened against 3CL-Pro and ACE2, respectively. Molecules with activities (AC50s) less than 50 μM and annotated PubChem ^45^ substance ids were collected. SMILES for each compound were retrieved using the PubChemPy API (https://pubchempy.readthedocs.io) and used to calculate cLogP values using RDkit-2019.09.3.0 (http://www.rdkit.org). Molecules with cLogP > 4 were drawn, visually inspected, and prioritized for testing. Finally, the SMILES of 2336 unique desalted molecules were selected from the SelleckChem library and were analyzed with Aggregation Advisor ^43^, a command line tool which calculates molecular similarity (Tanimoto Coefficients; Tc) between a list of molecules and a database of known aggregators (**SI Table 2**). Molecules with 1 > Tcs > 0.65 to a known aggregator and cLogP > 4 were drawn, visually inspected, and prioritized for testing. Percentages were calculated relative to the 2336 unique molecules in the library with identified SMILES.

### Compounds

All compounds are >95% pure by HPLC, as reported by the vendors. Compounds were ordered from Sigma Aldrich, Selleck Chem, Cayman Chemical or Medchem Express.

### Dynamic Light Scattering

To detect and quantify colloids, a DynaPro Plate Reader II (Wyatt Technologies) with a 60 mW laser at 830 nm wavelength and a detector angle of 158° was used; the beam size of the instrument was increased by the manufacturer to better enable detection of the colloids, which are larger than protein aggregates for which the instrument was designed. Samples were measured in 384 well plates with 30 μL loading and 10 acquisitions per sample. Compounds were dissolved in DMSO at 100x their final concentration and were diluted into filtered 50 mM KPi, pH 7.0, to obtain a final 1% DMSO concentration. Compounds were first tested at 3x the IC_50_ reported in the literature and if active further investigated in concentration-response. If no IC_50_ was available, compounds were tested at 100 μM. To calculate a CAC, each compound was serially diluted until substantial scattering disappeared; aggregating (>10^6^ scattering intensity) and non-aggregating (<10^6^ scattering intensity) portions of the data were fit with separate nonlinear regression curves, and the point of intersection was determined using GraphPad Prism software version 9.1.1 (San Diego, CA).

### Enzyme inhibition

MDH inhibition assays were performed at room temperature on a HP8453a spectrophotometer in kinetic mode using UV-Vis Chemstation software (Agilent Technologies) in methacrylate cuvettes (Fisher Scientific, 14955128) with a final volume of 1 mL for both control and test reactions. MDH (from Porcine Heart, 901643, Sigma-Millipore) was added to 50 mM KPi pH7 buffer for a final concentration of 2 nM. Compounds were dissolved in DMSO at 100x concentration, 10 μL of compound was used for a final DMSO concentration of 1%. After compound addition, the cuvette was mixed by pipetting up and down 5 times with a p1000, and the cuvette was then incubated for 5 minutes at room temperature. The reaction was initiated by the addition of 200 μM nicotinamide adenine dinucleotide (54839, Sigma Aldrich) and 200 μM oxaloacetic acid (324427, Sigma Aldrich) and the rate was monitored at 340 nm. A negative control was included in each run, in which 10 μL of DMSO without compound was added. The reactions were monitored for 90 seconds, and the initial rates were divided by the initial rate of the negative control to obtain the % inhibition and % enzyme activity. For dose response curves, 3 replicates were done for each concentration, the graphs were generated using GraphPad Prism version 9.1.1 (San Diego, CA).

### 3CL-Pro Kinetics Inhibition Assay

Fluorescent-quenched substrate with sequence rr-K(MCA)-ATLQAIAS-K(DNP)-COOH was synthesized via Fmoc solid-phase peptide synthesis as described.[Ravalin, 2019] Recombinant, active 3CL-Pro was expressed and purified as described.[Zhang 2020] Kinetic measurements were carried out in Corning black 384 well flat bottom plates and read on a BioTek H4 multimode plate reader. Quenched fluorogenic peptide had a final concentration of K_M_ = 10μM, and 3CL-Pro had a final concentration of 50nM. Reaction buffer was 20mM Tris, 150mM NaCl, 1mM EDTA, 0.05% Tween-20 (v/v), 1mM DTT, pH 7.4. Drugs were incubated with protease prior to substrate addition at 37°C for one hour. After incubation, substrate was added, and kinetic activity was monitored for one hour at 37°C. Initial velocities were calculated at 1 to 45 minutes in RFU/s. Velocities were corrected by subtracting the relative fluorescence of a substrate-only control and fraction activity was calculated using a substrate-corrected no-inhibitor control where DMSO was added instead of drug. Kinetics were carried out in triplicate.

## Associated Content

- Table of all compounds tested
- Figure of autocorrelation curves for all compounds considered aggregators
- Concentration response curves of the 17 literature compounds against 3CL-Pro in the presence of 0.05% Tween-20 Aggregate advisor analysis table

## Abbreviations Used

DLS: dynamic light scattering
Tc: Tanimoto Coefficients
CAC: critical aggregation concentration
MDH: malate dehydrogenase

## Acknowledgements

Supported by grants from the Defense Advanced Research Projects Agency HR0011-19-2-0020 and the NIH R35GM122481 (to BKS) and from NIAID (P50AI150476 to CSC). We thank Khanh Tang and John Irwin for help with Aggregation Advisor, and Isabella Glenn for help with aggregation assays.

## Notes

### Competing Interest Statement

The authors have declared no competing interest.

## References

1. Edwards, A. What Are the Odds of Finding a COVID-19 Drug from a Lab Repurposing Screen? J Chem Inf Model 2020, 60, 5727–5729.

2. Edwards, A.; Hartung, I. V. No shortcuts to SARS-CoV-2 antivirals. Science 2021, 373, 488–489.

3. Andreas Papapetropoulos, C. S. Inventing new therapies without reinventing the wheel: the power of drug repurposing. British journal of pharmacology 2018, 175,2, 165–167.

4. Dotolo, S.; Marabotti, A.; Facchiano, A.; Tagliaferri, R. A review on drug repurposing applicable to COVID-19. Brief Bioinform 2021, 22, 726–741.

5. Hossain, M. S.; Hami, I.; Sawrav, M. S. S.; Rabbi, M. F.; Saha, O.; Bahadur, N. M.; Rahaman, M. M. Drug Repurposing for Prevention and Treatment of COVID-19: A Clinical Landscape. Discoveries (Craiova) 2020, 8, e121.

6. Singh, T. U.; Parida, S.; Lingaraju, M. C.; Kesavan, M.; Kumar, D.; Singh, R. K. Drug repurposing approach to fight COVID-19. Pharmacol Rep 2020, 72, 1479–1508.

7. Saxena, A. Drug targets for COVID-19 therapeutics: Ongoing global efforts. J Biosci 2020, 45.

8. Baell, J. B.; Holloway, G. A. New substructure filters for removal of pan assay interference compounds (PAINS) from screening libraries and for their exclusion in bioassays. J Med Chem 2010, 53, 2719–40.

9. Baell, J. B. Feeling Nature’s PAINS: Natural Products, Natural Product Drugs, and Pan Assay Interference Compounds (PAINS). J Nat Prod 2016, 79, 616–28.

10. Dahlin, J. L.; Inglese, J.; Walters, M. A. Mitigating risk in academic preclinical drug discovery. Nat Rev Drug Discov 2015, 14, 279–94.

11. Thorne, N.; Auld, D. S.; Inglese, J. Apparent activity in high-throughput screening: origins of compound-dependent assay interference. Curr Opin Chem Biol 2010, 14, 315–24.

12. Lloyd, M. D. High-Throughput Screening for the Discovery of Enzyme Inhibitors. J Med Chem 2020, 63, 10742–10772.

13. Feng, B. Y.; Simeonov, A.; Jadhav, A.; Babaoglu, K.; Inglese, J.; Shoichet, B. K.; Austin, C. P. A high-throughput screen for aggregation-based inhibition in a large compound library. J Med Chem 2007, 50, 2385–90.

14. Jadhav, A.; Ferreira, R. S.; Klumpp, C.; Mott, B. T.; Austin, C. P.; Inglese, J.; Thomas, C. J.; Maloney, D. J.; Shoichet, B. K.; Simeonov, A. Quantitative analyses of aggregation, autofluorescence, and reactivity artifacts in a screen for inhibitors of a thiol protease. J Med Chem 2010, 53, 37–51.

15. Doak, A. K.; Wille, H.; Prusiner, S. B.; Shoichet, B. K. Colloid formation by drugs in simulated intestinal fluid. J Med Chem 2010, 53, 4259–65.

16. Seidler, J.; McGovern, S. L.; Doman, T. N.; Shoichet, B. K. Identification and prediction of promiscuous aggregating inhibitors among known drugs. J Med Chem 2003, 46, 4477–86.

17. Bojadzic, D.; Alcazar, O.; Buchwald, P. Methylene Blue Inhibits the SARS-CoV-2 Spike-ACE2 Protein-Protein Interaction-a Mechanism that can Contribute to its Antiviral Activity Against COVID-19. Front Pharmacol 2020, 11, 600372.

18. Ghahremanpour, M. M.; Tirado-Rives, J.; Deshmukh, M.; Ippolito, J. A.; Zhang, C. H.; Cabeza de Vaca, I.; Liosi, M. E.; Anderson, K. S.; Jorgensen, W. L. Identification of 14 Known Drugs as Inhibitors of the Main Protease of SARS-CoV-2. ACS Med Chem Lett 2020, 11, 2526–2533.

19. McGovern, S. L.; Shoichet, B. K. Kinase inhibitors: not just for kinases anymore. J Med Chem 2003, 46, 1478–83.

20. Ganesh, A. N.; Donders, E. N.; Shoichet, B. K.; Shoichet, M. S. Colloidal aggregation: from screening nuisance to formulation nuance. Nano Today 2018, 19, 188–200.

21. Blevitt, J. M.; Hack, M. D.; Herman, K. L.; Jackson, P. F.; Krawczuk, P. J.; Lebsack, A. D.; Liu, A. X.; Mirzadegan, T.; Nelen, M. I.; Patrick, A. N.; Steinbacher, S.; Milla, M. E.; Lumb, K. J. Structural Basis of Small-Molecule Aggregate Induced Inhibition of a Protein-Protein Interaction. J Med Chem 2017, 60, 3511–3517.

22. McGovern, S. L.; Helfand, B. T.; Feng, B.; Shoichet, B. K. A specific mechanism of nonspecific inhibition. J Med Chem 2003, 46, 4265–72.

23. Coan, K. E.; Maltby, D. A.; Burlingame, A. L.; Shoichet, B. K. Promiscuous aggregate-based inhibitors promote enzyme unfolding. J Med Chem 2009, 52, 2067–75.

24. Lak, P.; O’Donnell, H.; Du, X.; Jacobson, M. P.; Shoichet, B. K. A Crowding Barrier to Protein Inhibition in Colloidal Aggregates. J Med Chem 2021, 64, 4109–4116.

25. Feng, B. Y.; Toyama, B. H.; Wille, H.; Colby, D. W.; Collins, S. R.; May, B. C.; Prusiner, S. B.; Weissman, J.; Shoichet, B. K. Small-molecule aggregates inhibit amyloid polymerization. Nat Chem Biol 2008, 4, 197–9.

26. LaPlante, S. R.; Aubry, N.; Bolger, G.; Bonneau, P.; Carson, R.; Coulombe, R.; Sturino, C.; Beaulieu, P. L. Monitoring drug self-aggregation and potential for promiscuity in off-target in vitro pharmacology screens by a practical NMR strategy. J Med Chem 2013, 56, 7073–83.

27. Baker, J. D.; Uhrich, R. L.; Kraemer, G. C.; Love, J. E.; Kraemer, B. C. A drug repurposing screen identifies hepatitis C antivirals as inhibitors of the SARS-CoV2 main protease. PLoS One 2021, 16, e0245962.

28. Jin, Z.; Du, X.; Xu, Y.; Deng, Y.; Liu, M.; Zhao, Y.; Zhang, B.; Li, X.; Zhang, L.; Peng, C.; Duan, Y.; Yu, J.; Wang, L.; Yang, K.; Liu, F.; Jiang, R.; Yang, X.; You, T.; Liu, X.; Yang, X.; Bai, F.; Liu, H.; Liu, X.; Guddat, L. W.; Xu, W.; Xiao, G.; Qin, C.; Shi, Z.; Jiang, H.; Rao, Z.; Yang, H. Structure of M(pro) from SARS-CoV-2 and discovery of its inhibitors. Nature 2020, 582, 289–293.

29. Zhu, W.; Xu, M.; Chen, C. Z.; Guo, H.; Shen, M.; Hu, X.; Shinn, P.; Klumpp-Thomas, C.; Michael, S. G.; Zheng, W. Identification of SARS-CoV-2 3CL Protease Inhibitors by a Quantitative High-Throughput Screening. ACS Pharmacol Transl Sci 2020, 3, 1008–1016.

30. Olaleye, O. A.; Kaur, M.; Onyenaka, C.; Adebusuyi, T. Discovery of Clioquinol and analogues as novel inhibitors of Severe Acute Respiratory Syndrome Coronavirus 2 infection, ACE2 and ACE2 - Spike protein interaction in vitro. Heliyon 2021, 7, e06426.

31. Han, Y.; Duan, X.; Yang, L.; Nilsson-Payant, B. E.; Wang, P.; Duan, F.; Tang, X.; Yaron, T. M.; Zhang, T.; Uhl, S.; Bram, Y.; Richardson, C.; Zhu, J.; Zhao, Z.; Redmond, D.; Houghton, S.; Nguyen, D. T.; Xu, D.; Wang, X.; Jessurun, J.; Borczuk, A.; Huang, Y.; Johnson, J. L.; Liu, Y.; Xiang, J.; Wang, H.; Cantley, L. C.; tenOever, B. R.; Ho, D. D.; Pan, F. C.; Evans, T.; Chen, H. J.; Schwartz, R. E.; Chen, S. Identification of SARS-CoV-2 inhibitors using lung and colonic organoids. Nature 2021, 589, 270–275.

32. Tripathi, P. K.; Upadhyay, S.; Singh, M.; Raghavendhar, S.; Bhardwaj, M.; Sharma, P.; Patel, A. K. Screening and evaluation of approved drugs as inhibitors of main protease of SARS-CoV-2. Int J Biol Macromol 2020, 164, 2622–2631.

33. Fu, W.; Chen, Y.; Wang, K.; Hettinghouse, A.; Hu, W.; Wang, J. Q.; Lei, Z. N.; Chen, Z. S.; Stapleford, K. A.; Liu, C. J. Repurposing FDA-approved drugs for SARS-CoV-2 through an ELISA-based screening for the inhibition of RBD/ACE2 interaction. Protein Cell 2021, 12, 586–591.

34. Ge, S.; Wang, X.; Hou, Y.; Lv, Y.; Wang, C.; He, H. Repositioning of histamine H1 receptor antagonist: Doxepin inhibits viropexis of SARS-CoV-2 Spike pseudovirus by blocking ACE2. Eur J Pharmacol 2021, 896, 173897.

35. Lin, C.; Li, Y.; Zhang, Y.; Liu, Z.; Mu, X.; Gu, C.; Liu, J.; Li, Y.; Li, G.; Chen, J. Ceftazidime is a potential drug to inhibit SARS-CoV-2 infection in vitro by blocking spike protein-ACE2 interaction. Signal Transduct Target Ther 2021, 6, 198.

36. Jang, W. D.; Jeon, S.; Kim, S.; Lee, S. Y. Drugs repurposed for COVID-19 by virtual screening of 6,218 drugs and cell-based assay. Proc Natl Acad Sci U S A 2021, 118.

37. Brimacombe, K. R.; Zhao, T.; Eastman, R. T.; Hu, X.; Wang, K.; Backus, M.; Baljinnyam, B.; Chen, C. Z.; Chen, L.; Eicher, T.; Ferrer, M.; Fu, Y.; Gorshkov, K.; Guo, H.; Hanson, Q. M.; Itkin, Z.; Kales, S. C.; Klumpp-Thomas, C.; Lee, E. M.; Michael, S.; Mierzwa, T.; Patt, A.; Pradhan, M.; Renn, A.; Shinn, P.; Shrimp, J. H.; Viraktamath, A.; Wilson, K. M.; Xu, M.; Zakharov, A. V.; Zhu, W.; Zheng, W.; Simeonov, A.; Mathe, E. A.; Lo, D. C.; Hall, M. D.; Shen, M. An OpenData portal to share COVID-19 drug repurposing data in real time. bioRxiv 2020.

38. White, M. A.; Lin, W.; Cheng, X. Discovery of COVID-19 Inhibitors Targeting the SARS-CoV2 Nsp13 Helicase. bioRxiv 2020.

39. Shoichet, B. K. Interpreting steep dose-response curves in early inhibitor discovery. J Med Chem 2006, 49, 7274–7.

40. McGovern, S. L.; Caselli, E.; Grigorieff, N.; Shoichet, B. K. A common mechanism underlying promiscuous inhibitors from virtual and high-throughput screening. J Med Chem 2002, 45, 1712–1722.

41. McGovern, S. L.; Helfand, B.; Feng, B.; Shoichet, B. K. A Specific Mechanism for Non-Specific Inhibition. J. Med. Chem. 2003, 46, 4265–72.

42. Coan, K. E.; Shoichet, B. K. Stability and equilibria of promiscuous aggregates in high protein milieus. Mol Biosyst 2007, 3, 208–13.

43. Irwin, J. J.; Duan, D.; Torosyan, H.; Doak, A. K.; Ziebart, K. T.; Sterling, T.; Tumanian, G.; Shoichet, B. K. An Aggregation Advisor for Ligand Discovery. J Med Chem 2015, 58, 7076–87.

44. Zhang, C. H.; Spasov, K. A.; Reilly, R. A.; Hollander, K.; Stone, E. A.; Ippolito, J. A.; Liosi, M. E.; Deshmukh, M. G.; Tirado-Rives, J.; Zhang, S.; Liang, Z.; Miller, S. J.; Isaacs, F.; Lindenbach, B. D.; Anderson, K. S.; Jorgensen, W. L. Optimization of Triarylpyridinone Inhibitors of the Main Protease of SARS-CoV-2 to Low-Nanomolar Antiviral Potency. ACS Med Chem Lett 2021, 12, 1325–1332.

45. Kim, S.; Chen, J.; Cheng, T.; Gindulyte, A.; He, J.; He, S.; Li, Q.; Shoemaker, B. A.; Thiessen, P. A.; Yu, B.; Zaslavsky, L.; Zhang, J.; Bolton, E. E. PubChem in 2021: new data content and improved web interfaces. Nucleic Acids Res 2021, 49, D1388–D1395.

