## Supporting Information for "Colloidal aggregators in biochemical SARS-CoV-2 repurposing screens"

**Affiliations:**

**Contents**:

1. Table showing all compounds tested
2. DLS autocorrelation curves for 17 literature compounds and 6 AggregationAdvisor compounds
3. Concentration response curves of the 17 literature compounds against 3CL-Pro in the presence of 0.05% Tween-20.
4. Aggregator Advisor analysis table

**SI Table 1.**

| Name of Drug | Source | DLS results | MDH result | Reversed by detergent? | Aggregator? |
|---|---|---|---|---|---|
| Manidipine | Literature | Particles | Inhibited | Yes | Yes |
| Anthralin | Literature | Particles | Inhibited | Yes | Yes |
| Cinacalcet | Literature | Particles | Inhibited | Yes | Yes |
| lercanidipine | Literature | Particles | Inhibited | Yes | Yes |
| Shikonin | Literature | Particles | Inhibited | Yes | Yes |
| Gossypol | Literature | Particles | Inhibited | Yes | Yes |
| Clotrimazole / Mycelex | Literature | Particles | Inhibited | Yes | Yes |
| TTNPB | Literature | Particles | Inhibited | Yes | Yes |
| Clioquinol | Literature | Particles | Inhibited | Yes | Yes |
| 4E1RCat | Literature | Particles | Inhibited | Yes | Yes |
| YLF-466D | Literature | Particles | Inhibited | Yes | Yes |
| Aripiprazole | Literature | Particles | Inhibited | Yes | Yes |
| Hemin | Literature | Particles | Inhibited | Yes | Yes |
| hematein | Literature | Particles | Inhibited | Yes | Yes |
| myricetin | Literature | Particles | Inhibited | Yes | Yes |
| Emodin | Literature | Particles | Inhibited | Yes | Yes |
| Hypericin | Literature | Particles | Inhibited | Yes | Yes |
| Deltarasin | Literature | Particles | 50% inhibition at 100 uM | No | |
| Dienestrol | Literature | No Particles | 56% inhibition at 100uM | No | |
| Altrenogest | Literature | No Particles | slight inhibition at 100uM | No | |
| BMS309403 | Literature | No Particles | 50% inhibition at 100uM | restored to 15% inhibition | |
| Mycophenolic acid | Literature | No Particles | 8% inhibition at 100uM | No | |
| CAY-10581 | Literature | No Particles | 10% inhibition at 25uM | No | |
| Cepharanthine | Literature | Particles | Crashed out of buffer | Crashed out of buffer | |
| Proflavin | Literature | Particles | No Inhibition | N/A | |

| | | | | |
|---|---|---|---|---|
| TBB | Literature | No Particles | Inhibited | Restored to 65% inhibition |
| fascaplysin | Literature | Not fully dissolved? | Not fully dissolved? | Not fully dissolved? |
| Indocyanine-green | Literature | Particles | 48% inhibition | No |
| Evans-blue | Literature | No Particles | Inhibited | Yes |
| Tannic Acid (Gallotannin) | Literature | No Particles | Inhibited | No |
| Quinicrine | Literature | No Particles | 16% inhibition | No |
| Verteporfin | Literature | Particles | Slight inhibition | No |
| PYR-41 | Literature | Particles | Slight inhibition | No |
| Calcitriol | Literature | Particles | No Inhibition | N/A |
| Chinofon | Literature | No Particles | 50% inhibition at 100uM | Yes |
| theaflavin | Literature | No Particles | 80% inhibition | |
| Thioridazine | Literature | No Particles | Inhibited | |
| Teicoplanin | Literature | No Particles | No Inhibition | N/A |
| Terfenadine | Literature | No Particles | No Inhibition | N/A |
| efonidipine | Literature | No Particles | No Inhibition | N/A |
| DBeQ | Literature | No Particles | No Inhibition | N/A |
| Walrycin B | Literature | No Particles | No Inhibition | N/A |
| TDZD-8 | Literature | No Particles | No Inhibition | N/A |
| chlorpheniramine | Literature | No Particles | No Inhibition | N/A |
| doxylamine | Literature | No Particles | No Inhibition | N/A |
| doxepin | Literature | No Particles | No Inhibition | N/A |
| Methylene blue | Literature | No Particles | No Inhibition | N/A |
| Chlorpromazine | Literature | No Particles | No Inhibition | N/A |
| Perphenazine | Literature | No Particles | No Inhibition | N/A |
| Fluphenazine | Literature | No Particles | No Inhibition | N/A |
| Trifluoperazine | Literature | No Particles | No Inhibition | N/A |
| Promethazine | Literature | No Particles | No Inhibition | N/A |
| Tiapride | Literature | No Particles | No Inhibition | N/A |
| Quetiapine | Literature | No Particles | No Inhibition | N/A |
| Telaprevir | Literature | No Particles | No Inhibition | N/A |
| Blonanserin | Literature | No Particles | No Inhibition | N/A |

| | | | | | |
|---|---|---|---|---|---|
| Adapalene/Differin | Aggregator Advisor | Particles | Inhibited | Yes | Yes |
| Dracohodin perchlorate | Aggregator Advisor | Particles | Inhibited | Yes | Yes |
| Buparvaquone | Aggregator Advisor | Particles | Inhibited | Yes | Yes |
| Bifonazole | Aggregator Advisor | Particles | Inhibited | Yes | Yes |
| Alpha-Tochopherol | Aggregator Advisor | Particles | Inhibited | Yes | Yes |
| Bazedoxifene | Aggregator Advisor | Particles | Inhibited | Yes | Yes |
| Vitamin A | Aggregator Advisor | Particles | 15% inhibition | No | |
| Oxyclozanide | Aggregator Advisor | No Particles | 80% inhibition | Restored to 40% inhibition | |
| Tocofersolan | Aggregator Advisor | Particles | No Inhibition | N/A | |
| Clomifene | Aggregator Advisor | No Particles | 45% inhibition | No | |
| Flavoxate | Aggregator Advisor | No Particles | 15% inhibition | No | |
| Ipriflavone | Aggregator Advisor | Particles | Inconsistent | N/A | |
| Berbamine | Aggregator Advisor | No Particles | No Inhibition | N/A | |
| Candesartan | Aggregator Advisor | No Particles | No Inhibition | N/A | |
| Amodiaquine | Aggregator Advisor | No Particles | No Inhibition | N/A | |

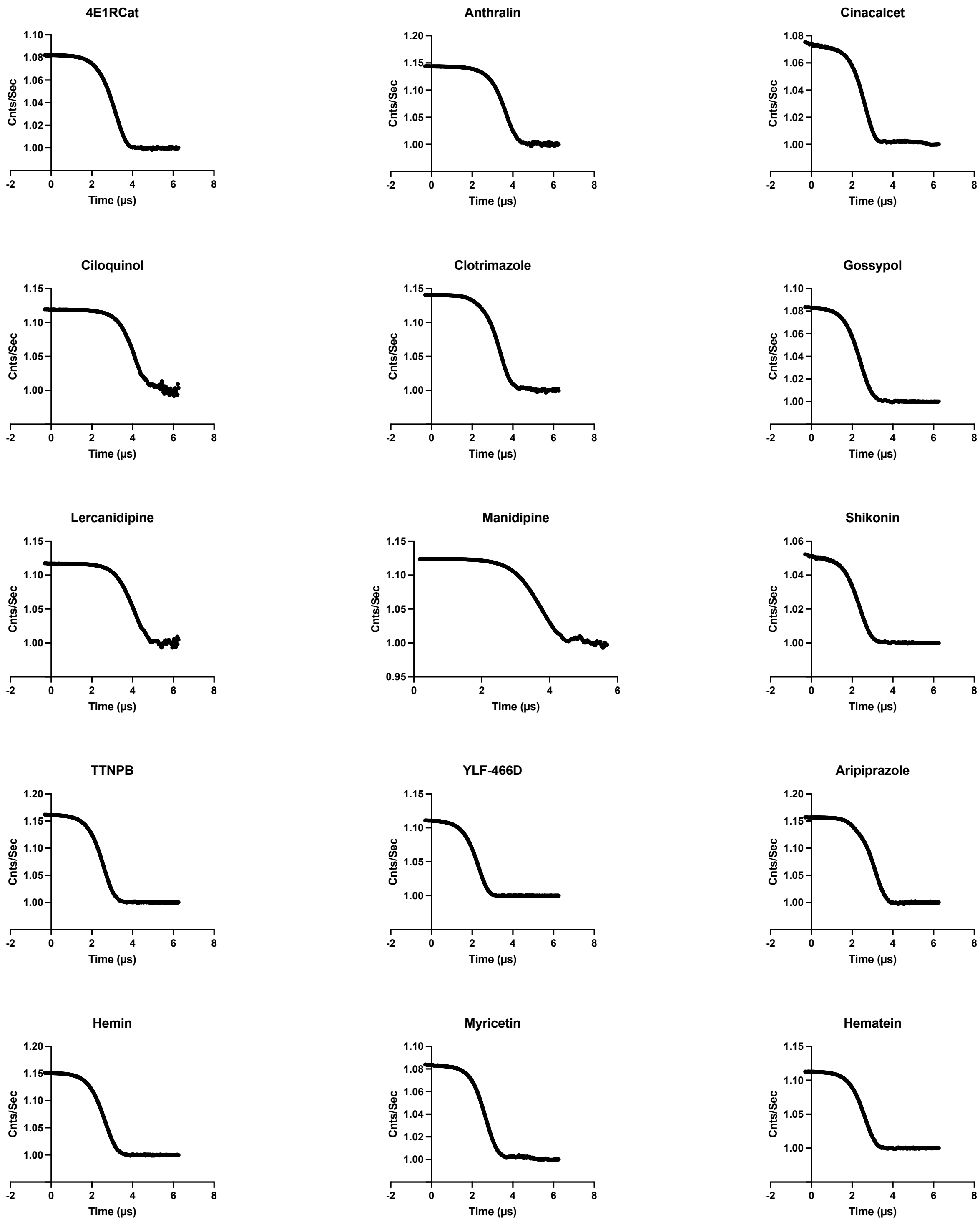

**Fig S1. DLS autocorrelation curves for literature reported hits.** Drug concentrations were at 3x $IC_{50}$ measured for MDH.

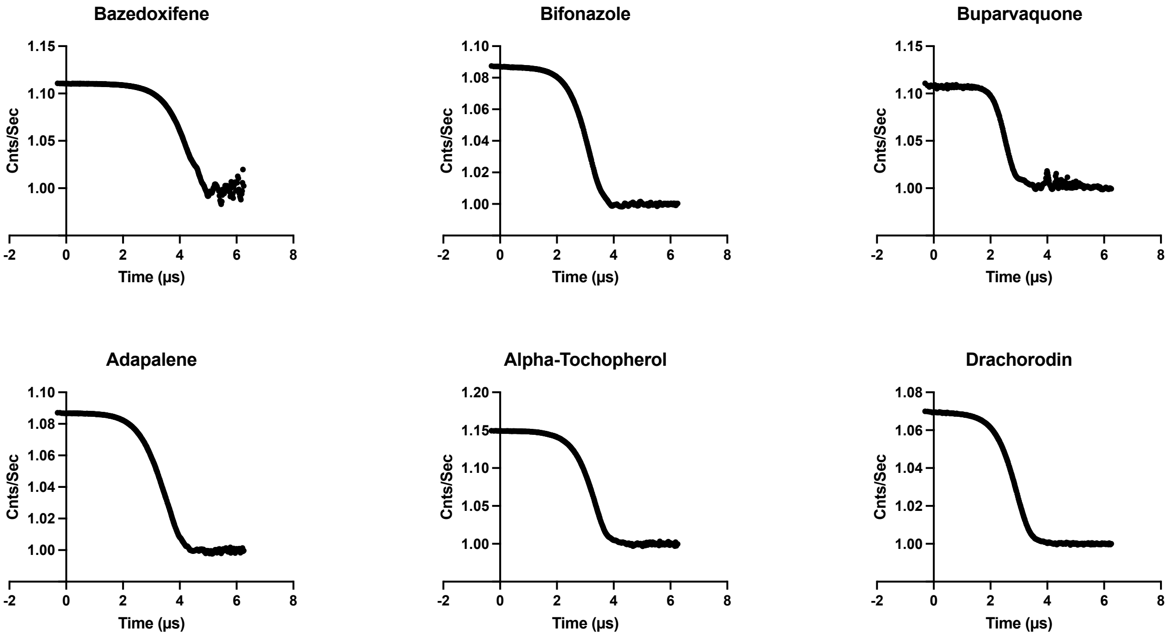

**Fig S2. DLS autocorrelation curves for drugs drawn from the repurposing library.** Drug concentrations were at 3x $IC_{50}$ measured for MDH.

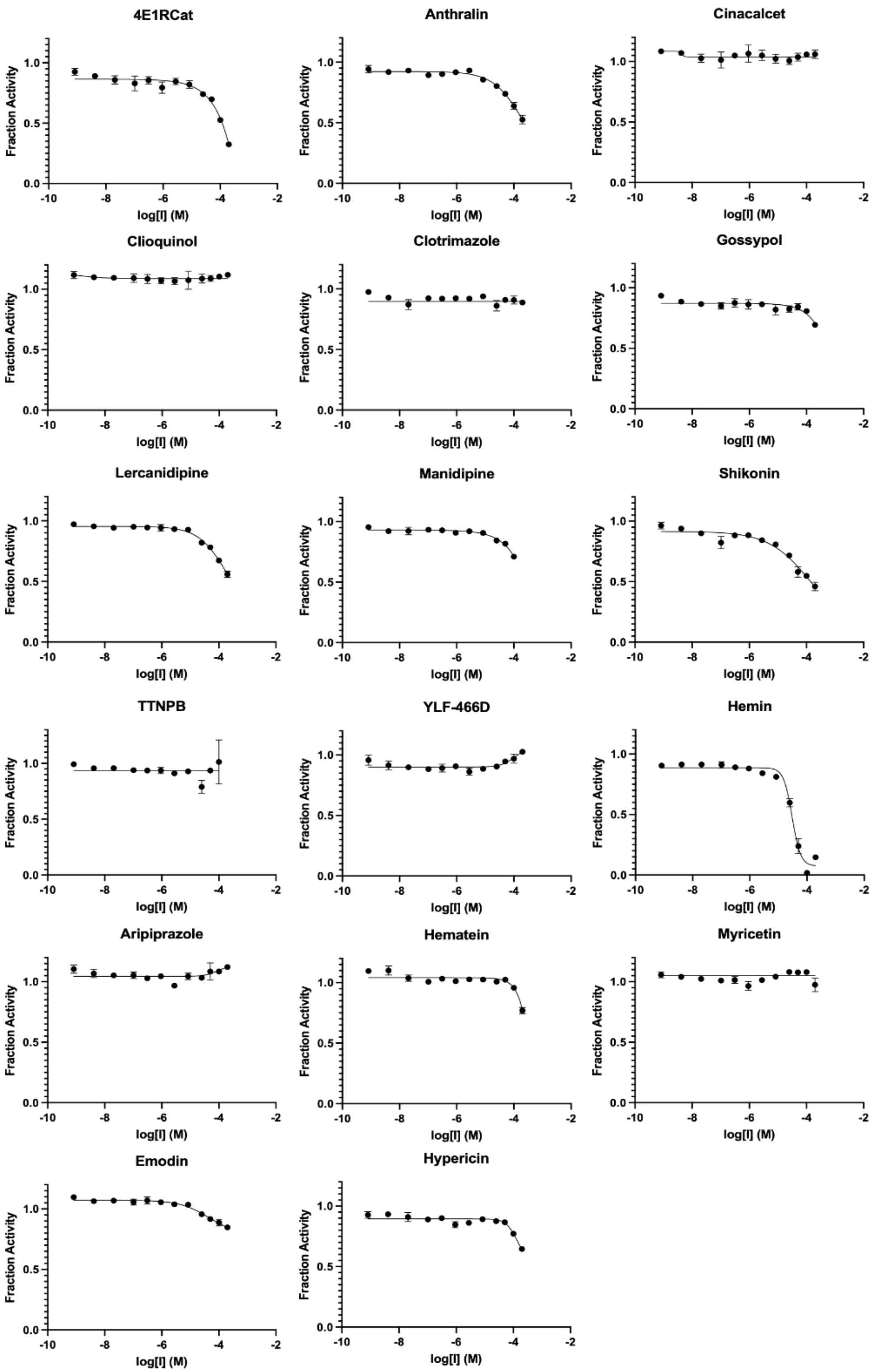

**Fig S3. Concentration response curves for literature compounds with 3CL-Pro in the presence of detergent.** With the exception of hemin, drugs at concentrations up to 200μM showed little potency on the enzyme in the presence of 0.05% Tween-20.

1. **SI Table 2.** Aggregator Advisor analysis table

| | TC | Name1 | Name2 |
|---|---|---|---|
| **0** | 0.83870965 | Butylated | hydroxytoluene |
| **1** | 1 | Triclabendazole | |
| **3** | 1 | Hydrocortisone | |
| **6** | 1 | Meclizine | 2HCl |
| **8** | 0.8115942 | (-)-Verbenone | |
| **9** | 0.8477366 | Indometacin | Sodium |
| **12** | 0.7075472 | Sodium_ferulate | |
| **13** | 1 | Gallamine | Triethiodide |
| **15** | 1 | Silymarin | |
| **20** | 1 | Methysergide | Maleate |
| **24** | 1 | Harmine | |
| **27** | 0.75090253 | ADP | |
| **28** | 0.88586956 | Bivalirudin | Trifluoroacetate |
| **29** | 0.7777778 | Norgestrel | |
| **31** | 0.7181818 | Levocetirizine | Dihydrochloride |
| **33** | 0.7916667 | Sparfloxacin | |
| **35** | 0.74864864 | Relugolix | |
| **49** | 1 | Sanguinarine | |
| **51** | 0.81526107 | Darunavir | |
| **52** | 0.8674699 | Lidocaine | |
| **53** | 0.79087454 | Danofloxacin | Mesylate |
| **55** | 0.7027027 | Bisoprolol | fumarate |
| **57** | 0.7637795 | Mestranol | |
| **59** | 1 | Clioquinol | |
| **63** | 0.9558824 | Diosmetin | |
| **112** | 0.7590361 | Norfloxacin | |
| **113** | 0.71428573 | Sulfalene(SMPZ) | |
| **114** | 0.78 | Cloxiquine | |
| **116** | 0.82978725 | Dextromethorphan | hydrobromide |
| **120** | 0.70434785 | Bevantolol | hydrochloride |

| | | | |
|---|---|---|---|
| **122** | 1 | Tinoridine | hydrochloride |
| **125** | 1 | Fulvestrant | |
| **127** | 0.7605634 | Benzocaine | |
| **129** | 1 | Sulconazole | Nitrate |
| **131** | 0.8535565 | uridine | triacetate |
| **133** | 0.97633135 | Fangchinoline | |
| **135** | 0.7848101 | Oleanolic | Acid |
| **137** | 0.75238097 | Chloroxine | |
| **139** | 0.89 | Dyclonine | HCl |
| **141** | 0.7777778 | Metaraminol | Bitartrate |
| **142** | 0.7615894 | α-Hederin | |
| **143** | 0.9508197 | Formononetin | |
| **145** | 0.7012195 | Tamibarotene | |
| **146** | 0.7248908 | Nilotinib | (AMN-107) |
| **150** | 0.7657993 | Epirubicin | HCl |
| **151** | 0.7735849 | Lathyrol | |
| **152** | 0.7773585 | Adenosine5'-monophosphate | monohydrate |
| **153** | 0.71875 | Caffeic | Acid |
| **155** | 0.784 | Promestriene | |
| **157** | 0.74703556 | Nicardipine | |
| **161** | 0.87401575 | Albendazole | |
| **163** | 0.76582277 | Primaquine | Diphosphate |
| **164** | 0.7225131 | Sertaconazole | nitrate |
| **168** | 1 | Embelin | |
| **172** | 0.9358974 | Pregnenolone | |
| **176** | 0.8055556 | Amiloride | HCl |
| **180** | 0.80124223 | Dipotassium | glycyrrhizinate |
| **181** | 0.95 | Isorhamnetin | |
| **228** | 0.83225805 | Melatonin | |
| **231** | 1 | Itraconazole | |
| **233** | 0.70711297 | UTP, | Trisodium |
| **234** | 0.71666664 | Teneligliptin | hydrobromide |
| **235** | 0.7058824 | Deferiprone | |

| | | | |
|---|---|---|---|
| 236 | 0.8695652 | Linoleic | acid |
| 237 | 0.875 | Ofloxacin | |
| 239 | 0.70989764 | Nadifloxacin | |
| 240 | 0.70652175 | Misoprostol | |
| 242 | 0.7619048 | Pirarubicin | |
| 243 | 0.7225807 | 5-Methoxytryptamine | |
| 245 | 0.8363636 | Tioconazole | |
| 249 | 0.8039216 | Salicylanilide | |
| 251 | 0.9970588 | Irinotecan | HCl |
| 252 | 0.7529412 | Inosine | |
| 253 | 0.7878788 | 8-Hydroxyquinoline | |
| 254 | 0.7222222 | Neohesperidin | |
| 255 | 0.7173913 | Alprostadil | |
| 257 | 0.7099237 | 2'-Deoxyguanosine | monohydrate |
| 258 | 0.7180617 | Vandetanib | (ZD6474) |
| 261 | 0.7297297 | Atenolol | |
| 271 | 0.7589286 | Droxidopa | |
| 274 | 0.77124184 | Sulfathiazole | |
| 276 | 0.8333333 | Pindolol | |
| 278 | 0.9020619 | Regorafenib | (BAY |
| 280 | 0.7647059 | Ethylparaben | |
| 282 | 0.7319149 | Afatinib | (BIBW2992) |
| 284 | 1 | Pranlukast | |
| 286 | 0.71532845 | Quinestrol | |
| 288 | 0.7069597 | Nelarabine | |
| 289 | 0.7848101 | Ursolic | Acid |
| 291 | 0.7746479 | 9-Aminoacridine | |
| 292 | 0.735426 | Ketanserin | |
| 293 | 0.7442623 | Dibutyryl-cAMP | (Bucladesine) |
| 294 | 0.73170733 | Flumequine | |
| 295 | 0.9109589 | Niclosamide | |
| 297 | 1 | Cinnarizine | |
| 300 | 0.73333335 | Estrone | |

| | | | |
|---|---|---|---|
| **302** | 0.7311828 | Tocofersolan | |
| **304** | 0.78039217 | Lomefloxacin | |
| **306** | 1 | Amiodarone | HCl |
| **310** | 0.7007874 | Mepivacaine | |
| **311** | 0.71634614 | Sulfamethoxypyridazine | |
| **313** | 0.708134 | Icotinib | |
| **315** | 0.7709677 | Arbidol | HCl |
| **318** | 0.74172187 | Ethopabate | |
| **319** | 0.71428573 | Cinepazide | maleate |
| **320** | 0.8671875 | Genistein | |
| **326** | 1 | Clopidogrel | |
| **328** | 1 | Cepharanthine | |
| **330** | 0.8034682 | Tetrahydropalmatine | |
| **332** | 0.7522124 | Carbenoxolone | Sodium |
| **333** | 0.9150943 | 4-Methylumbelliferone | (4-MU) |
| **348** | 0.9590164 | Diaveridine | |
| **350** | 0.73134327 | Evodiamine | |
| **353** | 0.70434785 | Levobetaxolol | HCl |
| **357** | 1 | Baicalein | |
| **406** | 0.75373137 | Oxibendazole | |
| **407** | 0.92134833 | 17-Hydroxyprogesterone | |
| **410** | 0.88659793 | Oxaprozin | |
| **412** | 1 | Raloxifene | |
| **419** | 0.8207547 | Deoxycorticosterone | acetate |
| **421** | 0.7886179 | 2-Methoxyestradiol | (2-MeOE2) |
| **427** | 0.720339 | Mevastatin | |
| **428** | 0.7058824 | Guaifenesin | |
| **429** | 0.8055556 | Testosterone | Enanthate |
| **432** | 0.7590361 | Pefloxacin | Mesylate |
| **433** | 0.84615386 | Emodin | |
| **434** | 0.720339 | Lovastatin | |
| **435** | 0.8448276 | Nonivamide | |
| **437** | 0.71910113 | Peimine | |

| | | | |
|---|---|---|---|
| 441 | 0.85365856 | L-Ornithine | |
| 442 | 0.9882353 | Phenazopyridine | HCl |
| 443 | 0.7027027 | Sitafloxacin | Hydrate |
| 444 | 0.86734694 | Prednisolone | |
| 445 | 0.7805907 | Alarelin | Acetate |
| 447 | 0.97633135 | Berbamine | |
| 449 | 1 | Clotrimazole | |
| 451 | 0.7777778 | Deferasirox | |
| 454 | 1 | Isotretinoin | |
| 455 | 0.88495576 | Fenofibric | acid |
| 457 | 0.71 | Marbofloxacin | |
| 458 | 0.7 | 3,4-Diaminopyridine | |
| 460 | 0.7473684 | Prilocaine | |
| 461 | 1 | Vemurafenib | (PLX4032, |
| 463 | 1 | Diiodohydroxyquinoline | |
| 467 | 0.7924528 | 3'-Fluoro-3'-deoxythymidine | (Alovudine) |
| 468 | 1 | Umbelliferone | |
| 474 | 0.71590906 | Madecassic | acid |
| 475 | 1 | Trimethoprim | |
| 477 | 0.8235294 | Osalmid | |
| 478 | 1 | Triamterene | |
| 480 | 0.7255814 | Cimifugin | |
| 481 | 1 | Indigo | |
| 483 | 0.8888889 | Methyldopa | |
| 486 | 1 | Bexarotene | |
| 490 | 0.8032129 | Enrofloxacin | |
| 492 | 0.80225986 | Cyclo(RGDyK) | |
| 493 | 0.78512394 | 2'-Deoxyadenosine | monohydrate |
| 494 | 0.74747473 | Proxyphylline | |
| 496 | 0.71153843 | Benorylate | |
| 497 | 1 | Econazole | |
| 501 | 0.7 | Tranilast | |
| 502 | 0.7118056 | Linagliptin | |

| | | | |
|---|---|---|---|
| **506** | 0.720339 | Simvastatin | |
| **507** | 1 | Quercetin | |
| **554** | 1 | Rilpivirine | |
| **562** | 0.90839696 | Ipriflavone | (Osteofix) |
| **567** | 1 | Fenofibrate | |
| **569** | 1 | Isoconazole | nitrate |
| **573** | 0.7805907 | Gonadorelin | Acetate |
| **575** | 0.8141026 | Ataluren | (PTC124) |
| **576** | 0.8888889 | Amprenavir | |
| **577** | 0.7805907 | Leuprolide | Acetate |
| **579** | 0.84864867 | CP21R7 | (CP21) |
| **585** | 0.78988326 | Fludarabine | |
| **586** | 0.74860334 | Tafamidis | |
| **588** | 0.89032257 | Sodium_Aescinate | |
| **589** | 0.7734375 | Piperine | |
| **590** | 0.7176471 | Hederagenin | |
| **591** | 0.7875 | D-Phenylalanine | |
| **592** | 0.75 | Canrenone | |
| **593** | 0.92134833 | Medrysone | |
| **596** | 0.75757575 | Guanfacine | Hydrochloride |
| **597** | 0.7006369 | Phenolphthalein | |
| **599** | 1 | Tretinoin | |
| **600** | 0.7894737 | Coumarin | |
| **601** | 1 | Sennoside | A |
| **603** | 0.75757575 | 1-Indanone | |
| **604** | 0.82474226 | Carbidopa | |
| **607** | 0.7366548 | Besifloxacin | HCl |
| **608** | 1 | Disulfiram | |
| **611** | 0.7293233 | Dracohodin | perochlorate |
| **617** | 0.902439 | L-Lysine | hydrochloride |
| **618** | 0.72602737 | (+)-(S)-Carvone | |
| **619** | 0.9240506 | Carboprost | |
| **621** | 0.78074867 | Thymopentin | |

| | | | |
|---|---|---|---|
| **624** | 0.72307694 | Difloxacin | hydrochloride |
| **625** | 0.7162162 | Rebamipide | |
| **626** | 1 | Fenbendazole | |
| **628** | 0.7389558 | Laquinimod | |
| **630** | 0.86734694 | Methylprednisolone | |
| **631** | 0.7777778 | Norethindrone | |
| **633** | 1 | Sofalcone | |
| **635** | 0.8142857 | Proflavine | |
| **636** | 0.7442748 | Yohimbine | HCl |
| **645** | 1 | Kaempferol | |
| **693** | 0.72602737 | Propylparaben | |
| **695** | 0.878327 | Tadalafil | |
| **721** | 1 | Sorafenib | |
| **723** | 0.7529412 | Isoprinosine | |
| **724** | 0.9652778 | Morin | |
| **771** | 1 | Nelfinavir | Mesylate |
| **773** | 0.7217391 | Dacomitinib | (PF299804, |
| **775** | 0.73333335 | L-Leucine | |
| **776** | 1 | Etravirine | (TMC125) |
| **784** | 0.84444445 | Eicosapentaenoic | Acid |
| **785** | 0.9705882 | Progesterone | |
| **787** | 0.7091837 | Etofylline | |
| **789** | 0.9758065 | Fenoldopam | mesylate |
| **790** | 0.74747473 | Posaconazole | |
| **792** | 0.92134833 | Medroxyprogesterone | |
| **795** | 0.8512397 | Indomethacin | |
| **798** | 0.9122807 | Casticin | |
| **844** | 0.71428573 | Dronedarone | |
| **849** | 0.7826087 | Oleic | Acid |
| **850** | 0.8090909 | Clomifene | citrate |
| **854** | 0.97633135 | (+)-Fangchinoline | |
| **856** | 0.7637795 | Zucapsaicin | |
| **857** | 0.8969072 | Flunarizine | 2HCl |

| | | | |
|---|---|---|---|
| **860** | 0.8309179 | Ruboxistaurin | (LY333531 |
| **866** | 0.7090909 | Stevioside | |
| **867** | 1 | Curcumin | |
| **872** | 0.75 | Lanatoside | C |
| **873** | 0.71428573 | Phenacetin | |
| **875** | 0.8372093 | Candesartan | |
| **879** | 0.7815126 | Nafarelin | Acetate |
| **881** | 0.8323699 | Cilengitide?trifluoroacetate | |
| **884** | 0.87068963 | 5,7-Dihydroxy-4-methylcoumarin | |
| **900** | 0.9240506 | Dinoprost | tromethamine |
| **902** | 0.7535545 | Sulfisoxazole | |
| **907** | 0.96276593 | Harmaline | |
| **909** | 0.8789809 | Fenticonazole | Nitrate |
| **913** | 0.8347107 | Vidarabine | |
| **914** | 0.74208146 | Doxifluridine | |
| **915** | 1 | Octocrylene | |
| **916** | 0.84126985 | Vitamin | A |
| **917** | 0.82954544 | Eriodictyol | |
| **918** | 0.78723407 | Glycyrrhetinic | acid |
| **922** | 0.92105263 | Adapalene | |
| **924** | 0.86470586 | Camptothecin | |
| **925** | 1 | Gatifloxacin | |
| **927** | 0.7848101 | Asiaticoside | |
| **928** | 0.73913044 | Prostaglandin | E2 |
| **930** | 0.88235295 | Flavone | |
| **975** | 0.7058824 | Ivermectin | |
| **976** | 0.9087591 | Moxifloxacin | |
| **978** | 0.75706214 | Cyclo | (-RGDfK) |
| **979** | 0.7 | Trans-Tranilast | |
| **980** | 0.8659794 | Cortisone | |
| **983** | 0.87068963 | Dihydrocapsaicin | |
| **984** | 0.7771429 | (+)-Catechin | |
| **988** | 0.7692308 | Hydroxyprogesterone | caproate |

| | | | |
|---|---|---|---|
| **991** | 0.7966102 | sulfaisodimidine | |
| **993** | 0.7619048 | Cyclizine | 2HCl |
| **998** | 0.75 | 4-Nitrophenol | |
| **999** | 0.74038464 | Actarit | |
| **1001** | 0.74626863 | Naftopidil | |
| **1002** | 1 | Gefitinib | (ZD1839) |
| **1004** | 0.8249027 | Terconazole | |
| **1006** | 0.7067669 | Eplerenone | |
| **1007** | 0.77619046 | Floxuridine | |
| **1008** | 0.7619048 | Meprednisone | |
| **1009** | 0.7619048 | Prednisone | |
| **1010** | 0.83707863 | Fluorescein | |
| **1012** | 0.84770113 | Rubitecan | |
| **1013** | 0.808 | Fosamprenavir | calcium |
| **1014** | 0.9074074 | AngiotensinII | human |
| **1016** | 1 | Miconazole | |
| **1020** | 0.7805907 | Leuprorelin | Acetate |
| **1022** | 1 | Tannic | acid |
| **1024** | 0.7094595 | Ranolazine | |
| **1025** | 0.70454544 | Quillaic | acid |
| **1026** | 0.9558824 | Sulfamethoxazole | |
| **1032** | 1 | Citropten | |
| **1037** | 1 | Silibinin | |
| **1042** | 0.8811594 | Belotecan | (CKD-602) |
| **1043** | 0.71597636 | Aspartame | |
| **1045** | 0.7222222 | 4-Aminoantipyrine | |
| **1048** | 0.85714287 | 2’-deoxyuridine | |
| **1049** | 0.91477275 | Taxifolin | (Dihydroquercetin) |
| **1056** | 0.8 | Ethinyl | Estradiol |
| **1058** | 0.9688716 | Balofloxacin | |
| **1060** | 1 | Clofoctol | |
| **1061** | 1 | Indirubin | |
| **1063** | 1 | Glafenine | HCl |

| | | | |
|---|---|---|---|
| **1064** | 0.7821782 | Etonogestrel | |
| **1065** | 0.8681948 | Topotecan | |
| **1066** | 0.80097085 | Idoxuridine | |
| **1067** | 0.7078652 | Saccharin | |
| **1069** | 0.787037 | Bergapten | |
| **1077** | 0.7032967 | Propantheline | bromide |
| **1078** | 1 | Clofazimine | |
| **1080** | 0.72307694 | Sarafloxacin | HCl |
| **1081** | 0.70289856 | Clofarabine | |
| **1082** | 1 | Chlorotrianisene | |
| **1086** | 0.7385159 | ATP | |
| **1087** | 0.7076923 | Esmolol | |
| **1095** | 0.7153846 | Istradefylline | |
| **1096** | 0.7876106 | Drospirenone | |
| **1097** | 0.71359223 | Erlotinib | |
| **1099** | 0.70247936 | Dexamethasone | (DHAP) |
| **1100** | 1 | 4-Aminopyridine | |
| **1102** | 1 | Crizotinib | (PF-02341066) |
| **1104** | 0.74380165 | Wedelolactone | |
| **1108** | 0.99236643 | Methylene | Blue |
| **1110** | 0.7075472 | Ferulic | Acid |
| **1111** | 0.80124223 | Glycyrrhizin | (Glycyrrhizic |
| **1112** | 0.9047619 | Oxfendazole | |
| **1114** | 0.73513514 | Farrerol | |
| **1115** | 0.71359223 | Nalidixic | acid |
| **1116** | 0.88529414 | (S)-10-Hydroxycamptothecin | |
| **1117** | 0.9222222 | Esculetin | |
| **1123** | 0.88980716 | Methylcobalamin | |
| **1125** | 1 | Apigenin | |
| **1174** | 0.7028302 | Xanthinol | Nicotinate |
| **1176** | 0.9358974 | Dehydroepiandrosterone | (DHEA) |
| **1180** | 0.7079646 | Ethacridine | lactate |
| **1182** | 0.7076023 | Tocopherol | |

| | | | |
|---|---|---|---|
| **1184** | 0.70989764 | Azilsartan | |
| **1188** | 0.70718235 | Rufinamide | |
| **1189** | 0.82325584 | Clevudine | |
| **1191** | 0.8156425 | Sulfaphenazole | |
| **1194** | 1 | Chlorhexidine?2HCl | |
| **1196** | 0.71900827 | norethisterone | enanthate |
| **1197** | 0.71428573 | Bisphenol | A |
| **1198** | 0.75889325 | Cladribine | |
| **1199** | 1 | Hexachlorophene | |
| **1201** | 0.77380955 | Tacrine | HCl |
| **1202** | 0.7096774 | Imperatorin | |
| **1204** | 0.84090906 | Sulfapyridine | |
| **1206** | 0.7519084 | Guanosine | |
| **1207** | 0.9027027 | (-)-Epicatechin | gallate |
| **1213** | 0.7529412 | Pramoxine | HCl |
| **1215** | 0.7327189 | Frovatriptan | Succinate |
| **1218** | 0.75 | (6–ε–Aminocaproic | acid |
| **1219** | 0.78350514 | Proguanil | |
| **1221** | 0.7790698 | Propranolol | HCl |
| **1223** | 0.70642203 | Aniracetam | |
| **1224** | 0.70731705 | Xipamide | |
| **1225** | 1 | 4-Aminophenol | |
| **1226** | 0.8032129 | Ciprofloxacin | hydrochloride |
| **1228** | 0.70731705 | Digoxin | |
| **1229** | 0.8730159 | Carazolol | |
| **1231** | 0.78125 | Itopride | hydrochloride |
| **1233** | 0.74538743 | Manidipine | |
| **1237** | 0.83208954 | Bosentan | Hydrate |
| **1239** | 0.7169811 | Benidipine | HCl |
| **1241** | 0.78640777 | Afloqualone | |
| **1248** | 0.825 | Uridine | |
| **1250** | 0.8034682 | Rotundine | |
| **1252** | 0.7637795 | Capsaicin(Vanilloid) | |

| | | | |
|---|---|---|---|
| **1253** | 0.72 | Urapidil | HCl |
| **1256** | 0.7821782 | Hydroxyzine | 2HCl |
| **1258** | 0.75757575 | Tribenzagan | Hydrochloride |
| **1260** | 0.7916667 | Zidovudine | |
| **1261** | 0.93650794 | Thymidine | |
| **1263** | 0.70212764 | (S)-Glutamic | acid |
| **1264** | 0.84615386 | Sinensetin | |
| **1310** | 0.8165939 | Roquinimex | |
| **1313** | 0.94382024 | Corticosterone | |
| **1316** | 0.8648649 | Estradiol | |
| **1320** | 0.7217391 | AKBA | |
| **1321** | 0.75336325 | SulfadiMethoxine | sodium |
| **1323** | 0.8451613 | Sulfamethizole | |
| **1325** | 0.79310346 | Acemetacin | |
| **1328** | 0.7134146 | Ouabain | |
| **1329** | 0.8181818 | Isofraxidin | |
| **1333** | 0.90163934 | Daidzein | |
| **1334** | 0.7657993 | Daunorubicin | HCl |
| **1335** | 1 | Imatinib | (STI571) |
| **1339** | 0.7777778 | Ethisterone | |
| **1341** | 0.81707317 | Vorinostat | (SAHA, |
| **1342** | 0.7898089 | Hederacoside | C |
| **1343** | 0.8142857 | Mitotane | |
| **1344** | 0.726087 | Bazedoxifene | Acetate |
| **1347** | 1 | Carvedilol | |
| **1349** | 0.8347107 | Adenosine | |
| **1350** | 0.7175926 | Hesperidin | |
| **1351** | 0.7389163 | Dyphylline | |
| **1353** | 0.75213677 | Gramine | |
| **1355** | 1 | Rosmarinic | acid |
| **1358** | 0.7116279 | Cytidine | |
| **1359** | 0.77037036 | Phenylpiracetam | |
| **1361** | 0.8979592 | Amodiaquine | |

| | | | |
|---|---|---|---|
| **1362** | 0.71428573 | Beta | Carotene |
| **1363** | 0.9117647 | Diphenylamine | Hydrochloride |
| **1366** | 1 | Luteolin | |
| **1415** | 0.75206614 | Bifonazole | |
| **1417** | 0.728 | Megestrol | Acetate |
| **1418** | 0.73255813 | Exemestane | |
| **1419** | 0.7368421 | Triclocarban | |
| **1420** | 0.8495575 | Estriol | |
| **1422** | 0.7692308 | Oxyclozanide | |
| **1423** | 1 | Escin | |
| **1424** | 0.72649574 | Talniflumate | |
| **1425** | 0.74157304 | Dichlorophene | |
| **1427** | 0.8516746 | Trifluridine | |
| **1429** | 0.72820514 | Oxytocin | (Syntocinon) |
| **1430** | 0.7008547 | Clofibrate | |
| **1432** | 0.70183486 | Masitinib | (AB1010) |
| **1433** | 0.88529414 | Hydroxy | Camptothecine |
| **1434** | 1 | Scopoletin | |
| **1440** | 0.70866144 | Trelagliptin | |
| **1441** | 0.9230769 | 7-Methoxycoumarin | |
| **1448** | 0.74285716 | Indobufen | |
| **1449** | 0.9166667 | Yangonin | |
| **1451** | 1 | Esculin | |
| **1455** | 0.85393256 | Daphnetin | |
| **1460** | 0.97633135 | Tetrandrine | |
| **1462** | 1 | Dithranol | |
| **1465** | 1 | Celecoxib | |
| **1467** | 0.7848101 | Madecassoside | |
| **1468** | 0.80841124 | Flavoxate | HCl |
| **1469** | 0.8309859 | Enzastaurin | (LY317615) |
| **1475** | 0.7631579 | Buparvaquone | |
| **1479** | 1 | Brivudine | |
| **1481** | 0.75 | isoleucine | |

| | | | |
|---|---|---|---|
| **1482** | 0.7263158 | Ademetionine | |
| **1483** | 0.7692308 | Metroprolol | succinate |
| **1495** | 0.7457627 | Ticlopidine | HCl |
